# Integrated multi-approaches reveal unique metabolic mechanisms of Vestimentifera to adapt to deep-sea hydrothermal vents

**DOI:** 10.1101/2023.06.25.546427

**Authors:** Qinglei Sun, Zihao Yuan, Yuanyuan Sun, Li Sun

**Affiliations:** CAS and Shandong Province Key Laboratory of Experimental Marine Biology, Center for Ocean Mega-Science, Institute of Oceanology, Chinese Academy of Sciences, Qingdao 266071, China; Laboratory for Marine Biology and Biotechnology, Laoshan Laboratory, Qingdao 266200, China; College of Earth and Planetary Sciences, University of Chinese Academy of Sciences, Beijing, China

**Author notes:** These two authors contributed equally to this work (Qinglei Sun, Zihao Yuan).

**Keywords:** Hydrothermal vents, Polychaeta, Siboglinidae, Vestimentiferan, Symbiosis, Comparative genomics, Trehaloneogenesis

## Abstract

Vestimentiferans (Siboglinidae, Polychaeta) thrive in deep-sea hydrothermal vents and depend on chemosynthetic symbiosis for nutrition. Currently, the central carbon metabolisms, especially the sugar synthesis pathways, of vestimentiferans remain obscure. In this study, the genome of the vestimentiferan *Arcovestia ivanovi* was obtained. Comparative genomics revealed that, unlike other Polychaeta, vestimentiferans possessed trehaloneogenesis and lacked gluconeogenesis. Transcriptome and metabolome detected the expression of trehalose-6-phosphate synthase (TPS), the key enzyme of trehaloneogenesis, and trehalose in vestimentiferan tissues, especially trophosome, suggesting the possibility of trehalose as the main blood sugar in vestimentiferans. Vestimentiferan TPS was most closely related to arthropod TPS and may be transferred from arthropods via transposons that existed in high densities around the vestimentiferan and arthropod TPS loci. Electron microscopy observed vestimentiferan symbionts with packed glycogen granules. Consistently, glycogen biosynthesis was present in vestimentiferan symbionts but absent in other Siboglinidae symbionts. Together this study revealed that vestimentiferans have evolved unique metabolic mechanisms to adapt to hydrothermal vents by utilizing trehaloneogenesis as the major sugar-synthesizing pathway, which produces trehalose to facilitate tolerance of the stresses (such as high temperature and H_2_S) of the vents. This study also indicated a critical role of bacterial glycogen biosynthesis in the highly efficient symbiont-vestimentiferan cooperation.

## Introduction

Since the discovery that the gutless giant tubeworm *Rifita pachyptila* of hydrothermal vents depended on chemosynthetic symbiosis for nutrition (1), deep-sea tubeworms, mainly Siboglinidae, have attracted a wide research attention across the world. In taxonomy, Siboglinidae was initially classified as the phyla Pogonophora and Vestimentifera, and later reclassified as members of Polychaeta, Annelida (2). To date, more than 200 species of siboglinids have been described, which belong to four main lineages, i.e., Vestimentifera, Sclerolinum (or Monilifera), Osedax, and Frenulata, all of which lack a gut and rely on bacterial symbionts for nutrition during adulthood (3, 4). Of these four groups of Siboglinidae, Vestimentifera is the most studied, due mainly to its ubiquitous population in the hydrothermal vents and cold seeps, which have been extensively studied in the past 40 years. At present, Vestimentifera consists of approximately 19 species, including *R. pachyptila*, *Arcovestia ivanovi*, *Tevnia jerichonana*, *Ridgeia piscesae*, *Lamellibrachia luymesi*, *Parescarpia echinospica*, *Oasisia alvinae*, *Seepiophila jonesi*, *Escarpia spicata*, and *Lamellibrachia columna* (5, 6). The adult body of vestimentiferan consists of the obturacular region, the vestimentum, the trunk, and the opisthosoma. The bacteriocyte-containing trophosome is in the trunk. Recently, several vestimentiferan genomes have been released, which increase our understanding of the adaptation mechanisms of vestimentiferans to symbiosis (7–11). For instance, certain genes thought to be involved in symbiont-host interaction appear to be expanded (7). There is also some evidence of horizontal gene transfer from bacteria to host in the vestimentiferan genomes (7).

Accumulating studies showed that the symbionts in Frenulate, Monilifera, and Vestimentifera were generally chemoautotrophic, whereas the symbionts in Osedax were heterotrophic (12). Currently, the vestimentiferan symbionts are the most studied, mainly in diversity and metabolism. The vestimentiferan symbionts are endosymbionts belonging to *Gammaproteobacteria* with sulfur-oxidizing potentials (6, 13). By now, no symbiont has been cultured, but more than 10 metagenome-based assembled genomes have been obtained (14–18), which revealed that vestimentiferan symbionts had similar carbon and energy metabolic potentials. All symbionts harbor the Calvin-Benson-Bassham (CBB) cycle and the reductive tricarboxylic acid cycle (15–17). Endosymbionts also have the pathways of glycolysis and tricarboxylic acid cycle (19). For energy metabolism, nitrate and oxygen possibly serve as electron acceptors, while hydrogen, sulfide, and thiosulfate are possible electron donors (7). The nutrient transport between symbiont−host has been studied in *R. pachyptila*, with some evidence suggesting host acquiring nutrients by symbiont digestion/lysis (20, 21), while other evidence suggesting that the symbiont provides the host with nutrients, such as succinate, by secretion (22, 23).

The central carbon metabolisms include glycolysis/glyconeogenesis, tricarboxylic acid cycle, and pentose phosphate pathway. In symbiosis, the central carbon metabolisms are a key to understanding the carbon flows in microbe, host, and between microbe and host. In vestimentiferan holobionts, the central carbon metabolisms of the host (vestimentiferans) remain to be investigated. In this study, the genome of *A. ivanovi* and the genomes of the symbionts of *A. ivanovi* and *L. columna* were obtained. Comparative genomics was conducted to examine the central carbon metabolisms in both vestimenferans and their symbionts. Furthermore, metatranscriptome, transcriptome, metabolome (to our best knowledge, for the first time), and transmission electron microscopy analyses were performed to better understand the metabolisms of the vestimentiferan holobionts. Our results revealed unique metabolic mechanisms of vestimentiferans and their symbionts, and add new insights into the evolution and environmental adaption of deep-sea Annelida.

## Materials and methods

### 16S rRNA gene amplicon sequencing

*A. ivanovi* and *L. columna* collected from hydrothermal vents were dissected into the plume, vestimentum, and trophosome. The trophosomes of nine *A. ivanovi* individuals and five *L. columna* individuals were used for the 16S rRNA gene amplicon analysis as reported previously (24) with some modifications. In brief, total DNA was extracted from the trophosome using the cetyltrimethylammonium bromide and sodium dodecyl sulfate method. The V3-V4 regions of the 16S rRNA gene were amplified as reported previously (25). Sequence data analyses were performed using QIIME2 and R packages (v3.2.0). The amplicon sequence variant (ASV) level alpha diversity indices were calculated using the ASV table in QIIME2 (26).

### Binning based on metagenome data

The trophosomes of one *A. ivanovi* individual and one *L. columna* individual were used to perform metagenomics. Total DNA was extracted and purified with the MetaHIT protocol as described previously (27). The DNA concentration was estimated by Qubit (Thermo Scientific, USA). The library was constructed and sequenced with the BGISEQ-500 protocol (28). Raw reads were filtered to remove low quality reads and host genomic DNA by SOAPnuke (29) and SOAPdenovo (30), respectively. Then the clean data were assembled with metaSPAdes (31), and open reading frame (ORF) was predicted with MetaGeneMark (32). All predicted genes were filter by 100 bp length cutoff, and were removed redundancies by cd-hit (33), resulting in a non-redundant gene catalogue. Gene profile was obtained as reported previously (27). MetaWRAP was used for binning MAG by default parameters (34). CheckM was used for assessing bins contamination and completeness (35).

### Metatranscriptomic analysis

The trophosomes of two *Arovestia* individuals and one *Lamellibrachia* individual were used for metatranscriptome sequencing. RNA was extracted with E.Z.N.A.® Soil RNA Kit (Omega bio-tek, USA). The cDNA library of each sample was constructed and sequenced on the HiSeq 3000 platform (Illumina, USA). Adapters and low-quality reads (base quality ≤ 20) were trimmed with Cutadapt. The clean reads were assembled and the genes were predicted with MEGAHIT (36) and MetaGeneMark (32). All predicted genes were filtered by 100 bp length cutoff, and were removed redundancies by cd-hit. Transcript expression levels were quantified in transcript per million (TPM) using kallisto (37). Metagenome gene catalog and metatranscriptome catalogue were searched against NR database, COG database, and KEGG database to obtain the taxonomy and function of the genes.

### Transmission electron microscopy (TEM)

For TEM analysis, the sample was fixed with 2.5% glutaraldehyde (Hushi, China), washed three times with 0.1 M PBS at 4 °C and then fixed with 1% osmium tetroxide (Ted Pella, USA) in PBS for 1.5 h, followed by washing as above. The sample was dehydrated in alcohol (Hushi, China), infiltrated with acetone (Tieta, China) and epoxy resin (SPI-CHEM, USA) mixture, embedded and polymerized in epoxy resin. Ultrathin sections were obtained with a Leica EM UC7 ultramicrotome (Leica Microsystems, Germany) and transferred onto copper grids covered with the formvar membrane (Electron Microscopy China, China). Uranyl acetate (2%) and lead citrate (Ted Pella INC, USA) were used for contrast staining. The sections were photographed with a transmission electron microscope (HT7700, Hitachi, Japan).

### Genome and transcriptome sequencing of *A. ivanovi*

Genomic DNA was isolated using the phenol-chloroform method. DNA degradation and contamination were monitored by electrophoresis in 1% agarose gels and pulsed-field gel electrophoresis. DNA purity was checked using a NanoPhotometer spectrophotometer (IMPLEN, USA). DNA concentration was measured using the Qubit DNA Assay Kit in Qubit 2.0 Flurometer (Life Technologies, USA). The partial mitochondrial cytochrome c oxidase I (COI) gene was amplified using the primer pairs COI L1490/COI H2198 (38) and sequenced. The sequences of other metazoans were collected from published data and Coarbitrator_COI_nuc.fa (39). The phylogenetic analysis was performed using IQ-TREE 2 with 1,000 bootstrap and best-fit model GTR+F+I+G4. Genome sequencing, assembly, annotation and transcriptome were performed in Novogene Bioinformatics Institute (Beijing, China). Both the Illumina Hiseq platform and the PacBio SEQUEL were used for the sequencing. For the Illumina Hiseq sequencing platform, 1.5 µg DNA per sample was used as the input material. Sequencing libraries were generated using NEB Next Ultra DNA Library Prep Kit (NEB, USA) following the manufacturer’s recommendations, and index codes were added to attribute sequences to each sample. The short but accurate reads from the Illumina platform were analyzed for genome survey and base level correction after the long-reads assembly. For the PacBio platform, genomic sequencing libraries were constructed according to the protocol recommended by PacBio. Long reads generated from the PacBio platform were used for genome assembly. RNA was extracted from the plume of six individuals using RNAiso Pure RNA Isolation Kit (Takara, Japan) and assessed for quality using NanoVue Plus spectrophotometer (GE Healthcare, USA). RNA-seq libraries were constructed as reported previously (40) and sequenced with Illumina HiSeq in paired-end 150 bp mode.

### De novo assembly of the *A. ivanovi* genome

The genome size of *A. ivanovi* was estimated using the Illumina sequencing data with the Kmer-based method (41). All subreads from the PacBio sequencing were assembled using the WTDBG software (https://github.com/ruanjue/wtdbg-1.2.8). The assembled sequence was polished using Quiver (SMRT Analysis v2.3.0) (42) with default parameters. To ensure the high accuracy of the genome assembly, several rounds of iterative error correction were performed using the Illumina clean reads. Subreads were assembled and error-corrected with WTDBG and Quiver. The completeness of the genome assembly was evaluated with Benchmarking Universal Single-Copy Orthologs (BUSCO) (43). The accuracy of the genome was evaluated using the Illumina short reads mapping with BWA (44). Additionally, the transcriptome reads were also mapped to the genome assembly using BLAT (45) with default parameters.

### Repetitive element annotation

The repetitive elements were identified via RepeatModeler 1.0.8 containing RECON (46) and RepeatScout with default parameters (47). The derived repetitive elements were searched against Repbase (48). The average number of substitution per site (K) for repeat unit was subtotaled. The K value was calculated based on the Jukes-Cantor formula: K=-300/4×Ln(1-D×4/300), with D representing the proportion of each repeat unit differing from the consensus sequences (Mouse Genome Sequencing Consortium, 2002).

### Protein-coding gene prediction and function annotation

Structural annotation of the genome incorporates homology-based prediction, ab initio prediction and RNA-Seq assisted prediction. For homogenous comparison prediction, the genome information of *Caenorhabditis elegans* (GCA_000002985.3), *Capitella teleta* (GCA_000328365.1), *Drosophila melanogaster* (GCA_000001215.4), *Helobdella robusta* (GCA_000326865.1), *Lingula anatine* (GCA_001039355.2), and *Octopus bimaculoides* (GCA_001194135.1) were obtained from the Ensembl database and NCBI database. The gene structures were predicted with TBLASTN (49) (E-value ≤ 1e−5) and GeneWise (50). For gene prediction based on Ab initio, Augustus (51), GeneID (52), GeneScan (53), GlimmerHMM (54) and SNAP (55) were used with optimized parameters.

Transcriptome reads assemblies were generated with Trinity for the genome annotation (56). To optimize the genome annotation, the RNA-Seq reads were aligned to genome sequence using Hisat2 (57) / TopHat (58) with default parameters to identify exon region and splice positions. The alignment results were then used as input for Stringtie (59)/ Cufflinks (60) with default parameters for genome-based transcript assembly. The non-redundant reference gene set was generated by merging genes predicted by three methods with EvidenceModeler using PASA (Program to Assemble Spliced Alignment) (61). Functional annotation of protein coding genes was evaluated with BLASTP (e-value 1e-05). The annotation information of the best BLAST hit, which was derived from database, was transferred to our gene set. Protein domains were annotated by searching SwissProt, InterPro, KEGG and Pfam databases.

### Gene expression and metabolomics analyses

The RNA-seq reads of tubeworms were processed and quality controlled via Trimmomatic (62). All trimmed reads were first mapped to the tubeworm genomes via HISAT2 (57) and HTSeq (63), and the final reads were processed with DESeq2 in iDEP (64) with FDR cutoff as 0.05 and minimum fold change as 2. The expression was plotted in Prism for visualization. Metabolomic analyses were performed with the trophosomes and vestimentums of three tubeworms as reported previously (65). Statistical significance was determined using unpaired Student’s t test.

### Data availability

The amplicon sequencing data were deposited in the Short Reads Archive (National Center for Biotechnology Information) under the accession number PRJNA834623. The 16S rRNA gene sequences of the *A. ivanovi* symbiont and the *L. columna* symbiont were deposited in GenBank under the accession numbers ON394528 and ON394527, respectively. The MAG genome sequences of the *A. ivanovi* symbiont and the *L. columna* symbiont were deposited in GenBank under the accession numbers PRJNA834915 and PRJNA834916, respectively. Metatranscriptomic sequences were deposited in the Short Reads Archive under the accession number PRJNA834913.

The COI gene sequences of *A. ivanovi* and *L. columna* were deposited in GenBank under the accession numbers MN385366 and OR140834, respectively. The raw reads of the transcriptome of *A. ivanovi* were deposited in GenBank under the accession number PRJNA985725. The assembled genome sequence of *A. ivanovi* was deposited in GenBank under the accession number PRJNA574016. The genomes and transcriptomes used for comparative analysis were collected from public databases: *Paraescarpia echinospica* (PRJNA625616), *Lamellibrachia luymesi* (PRJNA516467), *Capitella teleta* (PRJNA175705), *Dimorphilus gyrociliatus* (PRJEB37657), *Owenia fusiformis* (PRJEB38497), *Hydroides elegans* (PRJNA307290), *Paraescarpia echinospica* (PRJNA736170), *Lamellibrachia luymesi* (PRJNA516467). *Osedax frankpressi*, *Oasisia alvinae*, and *Riftia pachyptila*: https://github.com/ChemaMD/OsedaxGenome.

## Results

### Identification and characterization of hydrothermal vent vestimentiferans and their symbionts

The hydrothermal vent vestimentiferans *A. ivanovi* and *L. columna* were identified based on COI gene analysis, which showed that they shared high identities with *A. ivanovi* (99.40%) and *L. columna* (99.06%), respectively, and were phylogenetically clustered together with the *A. ivanovi* and *L. columna*, respectively, of Vestimentifera in the Siboglinidae family (**Figure 1A**). Since vestimentiferans rely on symbiotic bacteria for nutrition, we examined the symbiont community structures of *A. ivanovi* and *L. columna*. Fourteen 16S rRNA V3−V4 region amplicon libraries were constructed from the trophosomes of 9 *A. ivanovi* individuals and 5 *L. columna* individuals.

**Figure 1.**
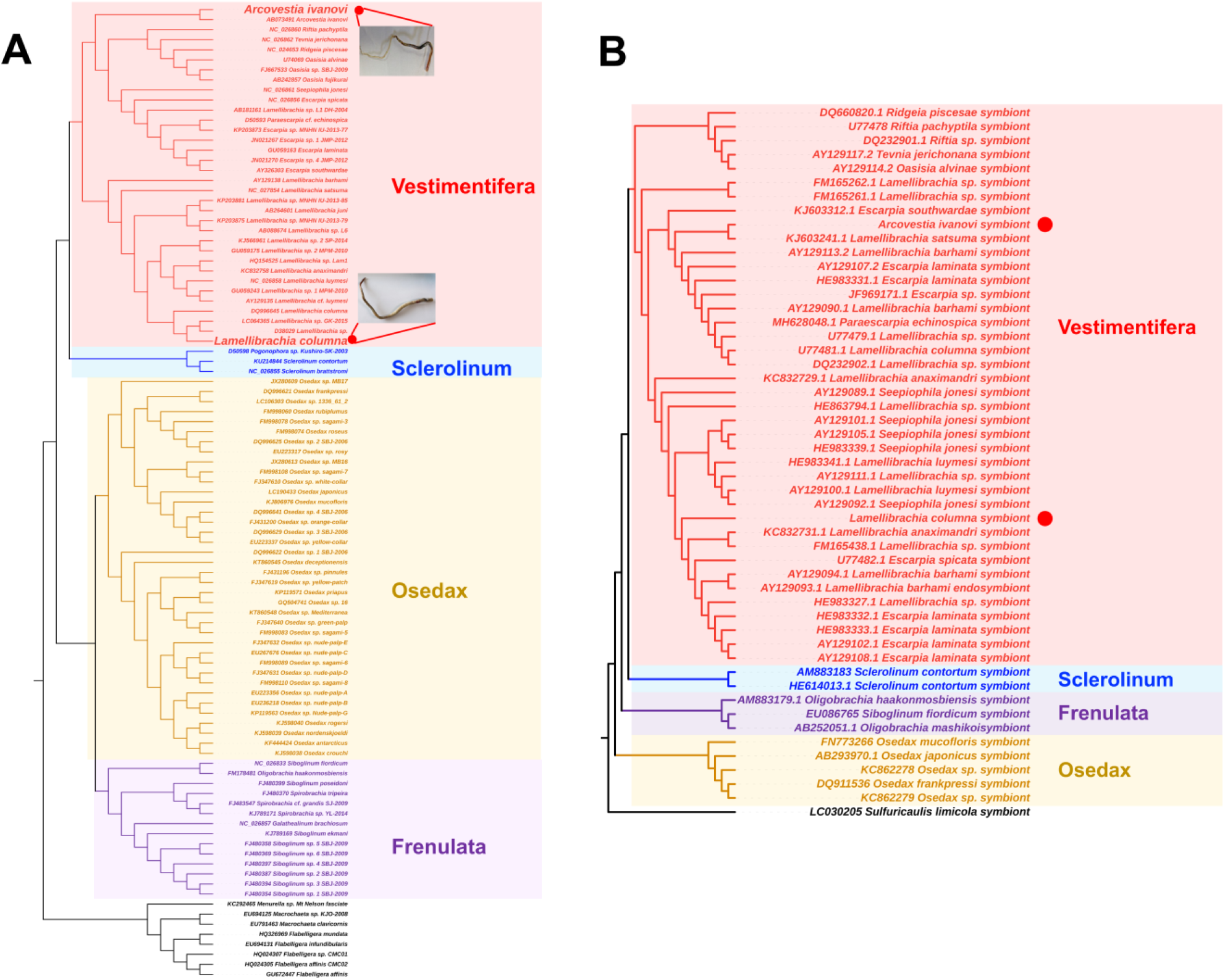
The phylogenetic analysis of siboglinids and their symbionts. (**A**) The COI gene-based phylogenetic tree of siboglinids constructed with iq-tree2. (**B**) The 16S rRNA gene-based phylogenetic tree of siboglinid symbionts constructed as above. The red dots in both panels indicate the *Arcovestia ivanovi* and *Lamellibrachia columna* from this study, and the photographs of these two species of tubeworms are shown in panel (A).

Community structure analysis showed that 98.5−99.8% and 59.4−87.7% tags of the *A. ivanovi* and *L. columna* symbiont libraries, respectively, were annotated to be of family Sedimenticolaceae (**Figure S1A**). For all 14 libraries, most tags were unclassified at the genus level, however, SUP 05 (0.2−19.1%) and *Sulfurovum* (1.0−13.8%) were relatively abundant in the *A. ivanovi* symbiont libraries (**Figure S1B**). All libraries were mainly composed of one ASV. ASV 42296 was most abundant in the *A. ivanovi* symbiont libraries, while ASV 24308 was most abundant in the *L. columna* symbiont libraries (**Figure S1C**). These results indicated that *Arcovestia* and *Lamellibrachia* each possessed mainly one ribotype of symbiont. The trophosomes of *A. ivanovi* and *L. columna* were each used to construct a metagenomic library. Only one nearly complete 16S rRNA gene (>1500 bp) of the symbiont was obtained from each library. The 16S rRNA gene sequences from the two libraries shared 98.4% identity and were clustered within the clade of vestimentiferan symbionts but segregated into different groups (**Figure 1B**). For both *A. ivanovi* and *L. columna*, only one high-quality metagenome-assembled genome (MAG) was obtained and used for further analysis.

### Metabolic features of the vestimentiferan symbionts revealed by meta-omics and electron microscopy

Genome analysis based on the above obtained MAGs indicated that both *A. ivanovi* and *L. columna* symbionts harbored all the genes of the CBB cycle (**Figure S1D**). Metatranscriptome analysis indicated that the key gene of the CBB cycle, ribulose-bisphosphate carboxylase, was expressed at a very high level in *A. ivanovi* and *L. columna* symbionts, with the TPM value ranging from 81.11 to 1986.90 (**Table S1**), suggesting that the CBB cycle was highly active. The *A. ivanovi* and *L. columna* symbionts also possessed all the genes of the rTCA cycle (**Figure S1D**), and ATP-dependent citrate lyase, the key gene of the rTCA cycle, was expressed at a high level (TPM of 14.06 to 25.28) (**Table S1**), suggesting that the rTCA cycle was active as well. In both *A. ivanovi* and *L. columna* symbionts, the gluconeogenesis pathway was not complete, and glucose-6-phosphatase, which transforms glucose-6-phosphate to glucose, was absent (**Figure S1D**).

However, the glycogen biosynthesis pathway was complete, and glucose-6-phosphate could be transformed into glycogen (**Figure S1D**). Metatranscriptome indicated that all the genes of glycogen biosynthesis were expressed, in particular glucose-1-phosphate adenylyltransferase, which was expressed at a high level (TPM of 15.51 to 26.28) in both *A. ivanovi* and *L. columna* symbionts (**Table S1**). Consistently, glycogen was detected in the holobiont by TEM and metabolome. TEM revealed lobule-like structures rich in bacteria in the trophosome (**Figure 2A**). Depending on their locations in the lobule, the symbiotic bacteria exhibited different morphologies. In the center of the lobule, the symbionts were small, with a diameter of 1−2 μm, and some bacteria were apparently in the stage of replication (**Figure 2A**). In the periphery, the symbionts were relatively large (2−4 μm in diameter) and filled with dark glycogen granules (**Figure 2A**). Some glycogen-packed cells appeared to undergo a process of lysis accompanying with glycogen release, and the cells eventually broke apart (**Figure 2B,C**). Comparative genomic analysis indicated that the glycogen biosynthesis pathway was present in most vestimentiferan symbionts but absent in Frenulate and Osedax symbionts (**Figure S1E**). In addition to the glycogen biosynthesis pathway, the complete riboflavin biosynthesis pathway was also present in the *A. ivanovi* and *L. columna* symbionts, and all the genes of this pathway were expressed (**Table S1**). Furthermore, the level of riboflavin in the trophosome was significantly and strikingly higher (1247 fold) than that in the vestimentum (**Table S2**).

**Figure 2.**
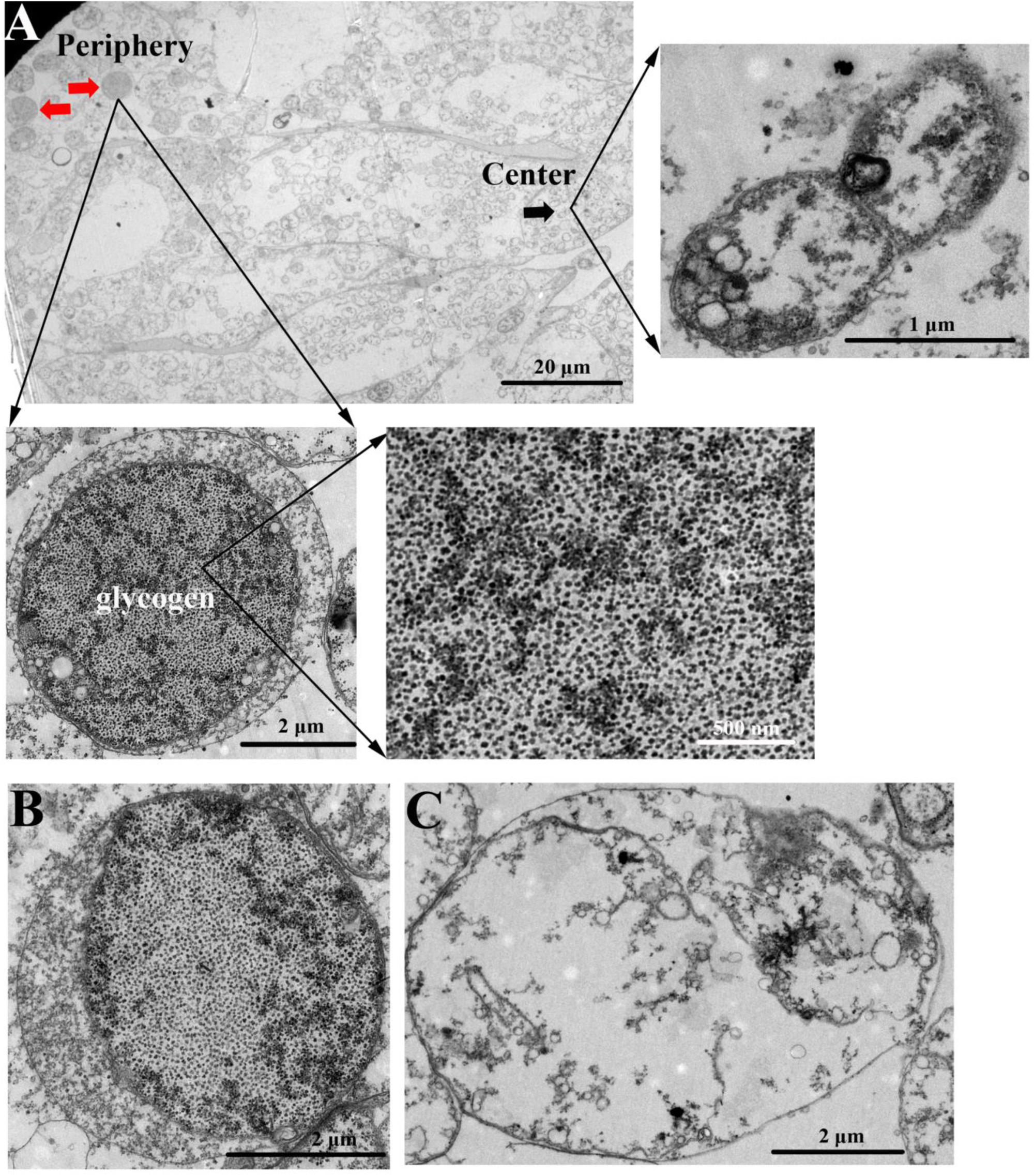
The symbionts in the *Lamellibrachia columna* trophosome observed with a transmission electron microscope. (**A**) The overall lobule-like structure. The black and red arrows indicate the smaller bacteria in the center and the larger bacteria in the periphery, respectively. The images of a dividing bacterium, a glycogen −filled bacterium, and the glycogen granules inside the bacterium are enlarged on the right, left bottom, and right bottom, respectively. (**B**) The symbiont in the process of releasing glycogen-containing cellular contents. (**C**) The representative of a symbiont with disintegrated structure after emptying glycogen contents.

### Assembly and annotation of the *A. ivanovi* genome

To explore the host-symbiont cooperation in metabolism, the genome of *A. ivanovi* was sequenced and assembled. Prior to assembly, the genome size of *A. ivanovi* was estimated to be approximately 853.7 Mb, with a heterozygous rate of 0.89%. The genome was assembled using a combination of Illumina paired-end and pacbio sequencing platforms (**Table S3**). The final assembled genome was 792.8 Mb and contained 9,469 contigs, with an N50 of 571.01 kb (**Table S4**). The completeness of the genome, evaluated using BUSCO against the eukaryotes genome, was 100% (249 complete, 6 fragmented, and no missing) (**Table S4**). The accuracy of the genome was evaluated using the Illumina short reads mapping with BWA. More than 98.1% of the reads were remapped to the assembled genome, thus validating the completeness of the genome. A total of 17,904 protein-coding genes were predicted in the genome, and 16,140 genes could be annotated using the InterPro, Swiss-Prot, NR and KEGG database (**Table S5**).

### The repeatome landscape of *A. ivanovi*

Since repetitive elements make up a large proportion of eukaryote genomes, we analyzed comprehensively the repeatomes of the tubeworms covering three families (**Table S6**). The *A. ivanovi* genome contained 477,439,658 bp (60.2% of the genome) repetitive element sequences, mainly class I transposable elements (124,047,905 bp), which accounted for 15.7% of the genome. The abundance of the class I transposable elements in *A. ivanovi* was comparable to that in other Siboglinidae species (14.4% in *Lamellibrachia luymesi* and 22.6% in *Paraescarpia echinospica*), but much higher than that in non-Siboglinidae species (5.6% in *Hydroides elegans* and 4.1% in *Owenia fusiformis*). Of the class I transposons, LINE expanded markedly in *A. ivanovi* (75.9% of the total transposable elements) as well as two other Siboglinidae species (71.3% and 61.7%, respectively, of the total transposable elements) and Serpulidae (59.5% of the total transposable elements), but made up only 36.2% of the total transposable elements in Oweniidae (**Figure 3A**). The Siboglinidae species were rich especially in LINE/CR1 and LINE/L2. Moreover, the LINE in *A. ivanovi* was not only abundant but also highly consensus, as reflected by the low substitution rates (**Figure 3B**).

**Figure 3.**
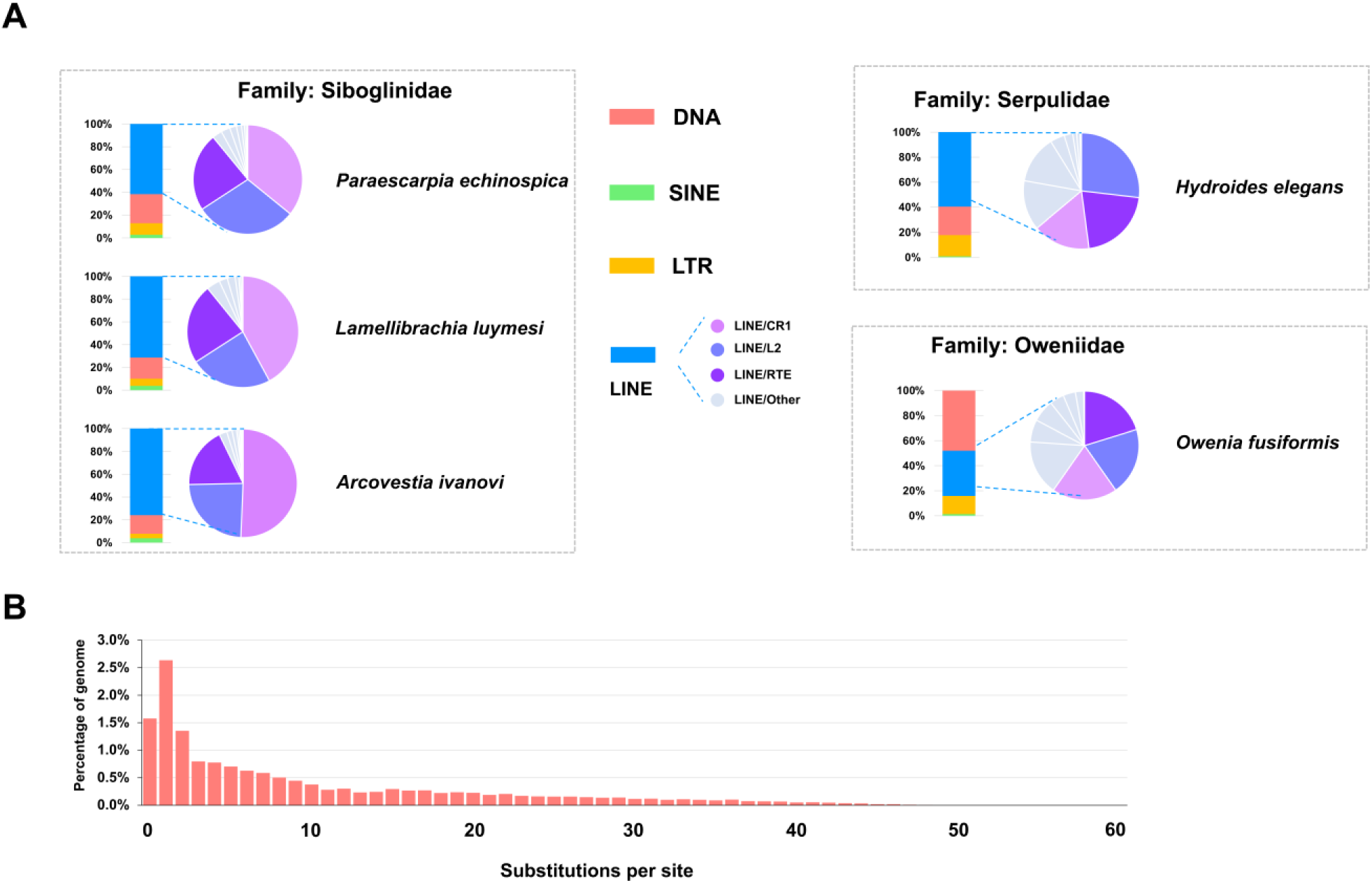
Analysis of the repetitive elements in tubeworms. (**A**) The abundance of the repetitive elements in Siboglinidae compared with that in Serpulidae and Oweniidae. (**B**) The divergence of LINE/CR1 in *Arcovestia ivanovi.* The y-axis represents the percentage of the genome comprised of repeat classes (%) and the x-axis represents the substitution rate from consensus sequences (%).

### The unique metabolism pathways in vestimentiferans

Currently, little is known about the central carbon metabolisms in vestimentiferans. To explore the carbon metabolism pathways in vestimentiferans, comparative genomics was conducted, which showed that the glycolysis pathway, tricarboxylic acid cycle, and pentose phosphate pathway were complete in all analyzed Polycheata genomes (**Table S7**). However, the sugar-synthesizing pathways differed markedly between symbiotic and non-symbiotic Polychaeta, as well as between different groups of Siboglinidae. All non-symbiotic Polychaeta possessed complete gluconeogenesis pathway but lacked trehalose 6-phophate synthase (TPS), the key enzyme for trehalose synthesis, and thus were probably unable to synthesize trehalose (**Figure 4A**). On the contrary, all vestimentiferans lacked glucose-6-phosphatase, the key enzyme for glucose synthesis, but possessed TPS, and therefore were able to synthesize trehalose (**Figure 4A,B**). Surprisingly, Osedax, another member of Siboglinidae, possessed neither TPS nor glucose-6-phosphatase (**Figure 4A**). Transcriptome analysis indicated that TPS was expressed in vestimentiferan tissues (**Figure S2**). In agreement, metabolome detected trehalose in vestimentiferan trophosome and vestimentum, especially the former (**Table S2; Figure 4C**). In contrast, although glucose is an important source of energy, it was not detected in the vestimentiferan metabolome. Sequence analysis showed that the TPS of five Siboglinidae species shared high similarities with each other (87.8−96.1%) and with the TPS of Arthropoda (58.5−73.1% in the top 10 alignments), but shared low similarities with the TPS of Nematoda (20.1−34.1%). Consistently, phylogenetic analysis showed that Annelida TPS was most closely related to Arthropoda TPS (**Figure 4D**). Furthermore, like many arthropod TPS, vestimentiferan TPS was a fused protein with both TPS/OtsA and TPP/OtsB domains (**Figure 4E)**. High densities of LINE repetitive elements with low substitution rates were found around the TPS loci of *A. ivanovi* and American lobster *Homarus american* (Arthropoda) (**Figure 4F**).

**Figure 4.**
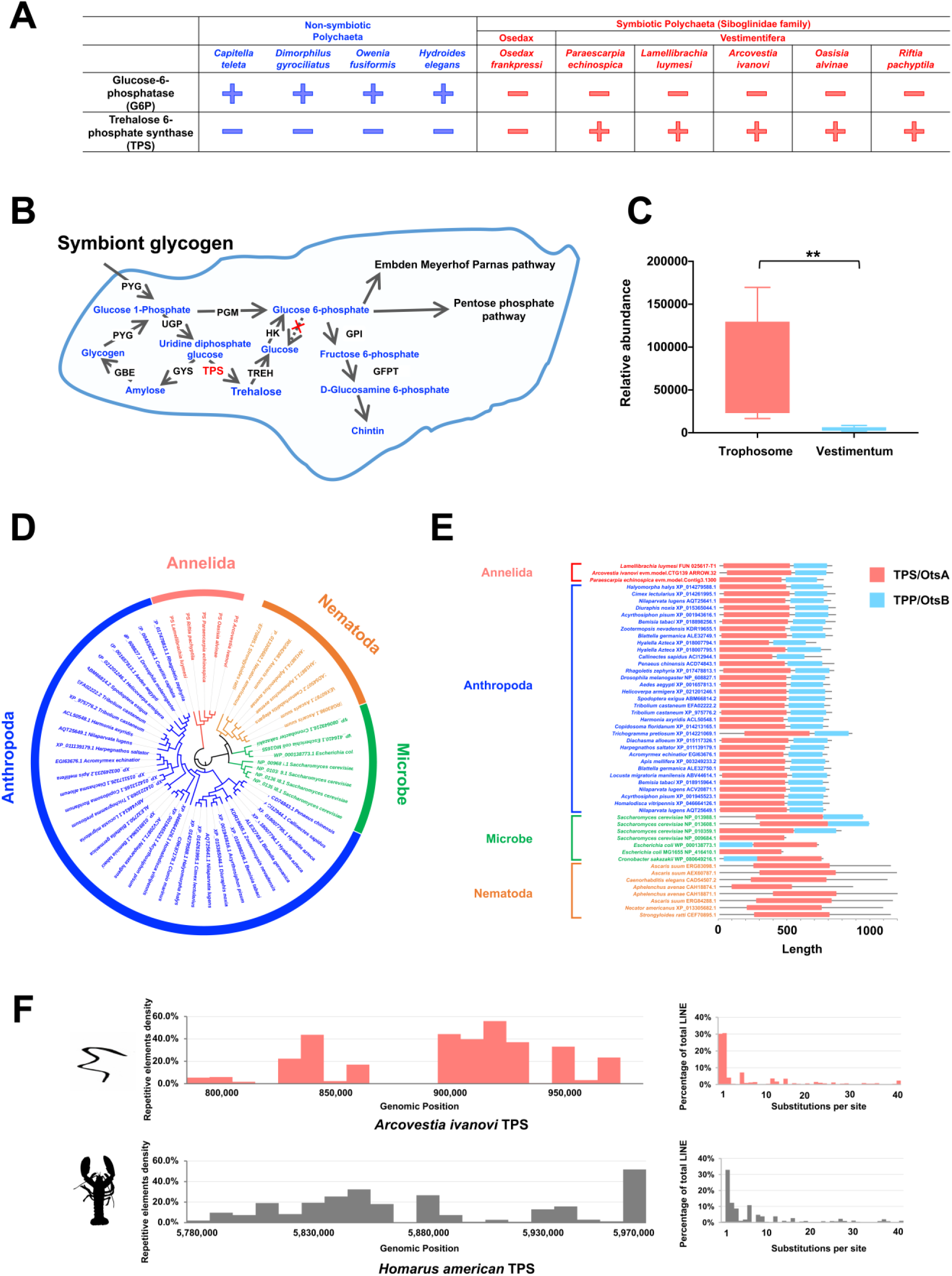
The trehalose biosynthesis pathway in Vestimentifera. (**A**) The distribution of trehalose 6-phosphate synthase (TPS) and glucose-6-phosphatase (G6P) in symbiotic and non-symbiotic Polychaeta. (**B**) The proposed carbon flow in Vestimentifera. The abbreviations of the enzymes are as follows: PYG: Glycogen phosphorylase; UGP: UTP-glucose-1-phosphate uridylyltransferase; GYS: Glycogen synthase; GBE: 1,4-alpha-glucan-branching enzyme; PGM: Phosphoglucomutase; TREH: Trehalase; HK: Hexokinase; GPI: Glucose-6-phosphate isomerase; GFPT1: Glutamine--fructose-6-phosphate aminotransferase. (**C**) The relative abundance of trehalose in vestimentiferan tissues detected by metabolome **, p<0.01. (**D**) The phylogenetic analysis of the TPS from Annelida, Arthropoda, Nematoda, and microbes. **E.** The comparison of the conserved domains of TPS from Annelida, Arthropoda, Nematoda, and microbes. **F.** The comparison of the LINE hot spots in *Arcovestia ivanovi* and American lobster *Homarus american*.

## Discussion

Vestimentiferans in hydrothermal vents can grow at an exceedingly high rate, for example, > 85 cm and > 30 cm increase in tube length per year for *R. pachyptila* and *T. jerichonana*, respectively (66). Vestimentiferans form very dense communities and accumulate a large amount of biomass, suggesting that the vestimentiferan-symbiont partnership is a highly successful strategy for adaptation to the vent surroundings. Since the symbionts provide the host vestimentiferans with abundant carbon compounds, the carbon metabolism of the symbionts plays an important role in the holobiont system. Glycogen, an essential carbon reserve for many organisms, has been detected in vestimentiferan symbionts and hosts via TEM and chemical analysis (21, 67). However, the glycogen biosynthesis pathway at the molecular level remains to be investigated. In this study, genome analysis indicated that the glycogen biosynthesis pathway was conserved in most vestimentiferan symbionts, but absent in Frenulate and Osedax symbionts. The presence of glycogen biosynthesis in vestimentiferans was verified by transcriptome, metabolome, and TEM analyses. By TEM, we found that the symbionts located in the lobule periphery accumulated much more glycogen than the symbionts in the lobule center, which was similar to the observation in *R.pachyptila*, another Vestimentifera member (67, 68). In our study, broken bacterial cells and glycogen release were observed in the periphery of the lobule. These results suggested that glycogen was an important carbon reserve in vestimentiferan symbionts and, after lysis/digestion of the symbionts, could be supplied to the host as a vital nutrient. Hence, glycogen biosynthesis is likely critical for the establishment and maintenance of the highly efficient symbiont-vestimentiferan cooperation. It is interesting that, in addition to glycogen biosynthesis, this study also found that riboflavin biosynthesis was active in both *A. ivanovi* and *L. columna*. Consistently, riboflavin was detected for the first time in Vestimentifera. Riboflavin, also called vitamin B2, is essential for all organisms, however, it is only known to be produced by plants and microbes. Its detection in Vestimentifera suggested that, in addition to glycogen, riboflavin may be another important nutrient supplied by the symbionts to the host vestimentiferans. In a previous report, riboflavin was shown to stimulate metamorphosis in the larvae of polychaete *Capitella telet*a (69). It is possible that the riboflavin produced by the vestimentiferan symbionts observed in our study may be involved in the metamorphosis, growth, and development of the host tubeworms.

Since sugar synthesis is critical for the survival of all animals, especially under conditions of nutrient deficiency, sugar-synthesizing pathways are expected to exist in all metazoan species (70). Gluconeogenesis is important for mammals and fish, because glucose, the product of this pathway, is utilized as the blood sugar (70). In contrast, trehaloneogenesis is important for many insects, because trehalose (a non-reducing disaccharide comprising two glucose molecules), the product of trehaloneogenesis, is the principal hemolymph sugar of insects (71). In some insects, such as the fruit fly, both trehaloneogenesis and gluconeogenesis are present, with the latter participating in neuronal signaling (72). To date, the sugar-synthesizing pathways in Annelida are largely unknown. In this study, the sugar-synthesizing pathways in Polychaeta were systematically analyzed via comparative genomics. Unexpectedly, we found that unlike non-symbiotic Polychaeta, vestimentiferans possessed only trehaloneogenesis and lacked gluconeogenesis, suggesting that trehalose, rather than glucose, was likely a vital energy source for vestimentiferans. Consistently, trehalose was detected in vestimentiferan organs, especially the trophosome. Since trophosome is the site of hematopoiesis and rich in blood vessels (5, 10), trehalose may be the principle blood sugar of vestementiferans. Trehalose is also known as a cryoprotectant and a heat bioprotectant for cells and proteins (73), and this property may enable trehalose to confer a surviving advantage upon vestementiferans. Previous reports showed that the vestimentiferan Riftia grew optimally at 25 ℃ but could not tolerate prolonged incubation in the higher temperature of 32 to 35°C (74). Since in hydrothermal vents, the distribution of H_2_S tends to be more abundant at the regions closer to the vent (and hence higher temperature), vestimentiferans may be exposed to lethal high temperatures in order to obtain the sulfide required to sustain the chemoautotrophic metabolism of their symbionts (74). Therefore, trehalose, as a heat protectant, may function to shield the vestimentiferans from the harm of the high temperature in the vents. In addition, trehalose was reported to participate in H_2_S resistance in maize (75). It is possible that, besides its heat-resistance property, trehalose may also play a role in the sulfide tolerance of vestimentiferans. Together, these results supported trehaloneogenesis as a critical mechanism for vestimentiferans to adapt to the specific conditions of hydrothermal vents.

In this study, TPS was detected only in vestimentiferans and not in other annelids, which raised the question of the origin of the vestimentiferan TPS. We found that the vestimentiferan TPS shared high levels of sequence similarities with arthropod TPS, suggesting the possibility of horizontal transfer of the TPS gene from Arthropada to Annelida. To date, very few horizontal gene transfer events have been verified in animals. Transposable element is a propelling force for genomic evolution, and horizontal transfers of transposons in eukaryotes have been reported (76, 77). By comparative analysis, we indeed observed a high abundance of transposable elements surrounding the TPS loci of *A. ivanovi* and the arthropod *H. American* (lobster), suggesting active genomic activity adjacent to the TPS genes in these animals. Given that some deep-sea arthropods, such as *Alvinocaris longirostris* (shrimp) and *Shinkaia crosnieri* (crab) (24, 78), inhabit deep-sea hydrothermal vents/cold seeps like vestementiferans and reproduce via external fertilization, horizontal gene transfer between these arthropods and annelids (vestementiferans) mediated by transposable elements is possible and might be one of the causative forces that have led to TPS transfer across species. The acquisition of TPS enables vestementiferans to synthesize trehalose. As discussed above, trehalose is likely the primary energy source of vestementiferans and may facilitate the resistance of vestementiferans against the high temperature and H_2_S in hydrothermal vents. In addition, in insects, trehalose is an essential substrate for chitin synthesis, which is directly affected by TPS (79). Like insects, vestementiferans, which are characterized by conspicuous chitinous tubes, require chitin synthesis. Hence, the acquisition of TPS probably endows vestimentiferans with a growth and surviving advantage that is conducive to the flourish of vestimentiferans in deep-sea hydrothermal vents.

It is interesting that in this study, we found that the sugar-synthesizing pathways in Siboglinidae differed markedly among different groups. While Vestimentifera possessed trehalogenensis and lacked gluconeogenesis, Osedax possessed neither gluconeogenesis nor trehalogenensis. Since no genome sequence of Frenulate or Monilifera has been reported, the sugar-synthesizing pathways in these two groups are unclear. Previous phylogenetic analyses showed that Vestimentifera emerged most recently in Siboglinidae, while Frenulate formed the basal group of this family, and Osedax was positioned between Frenulate and Vestimentifera (4, 80, 81).

Combining these observations with our results, it can be hypothesized that the ancestor siboglinida, like other groups of Polychaeta, possessed the complete pathway of gluconeogenesis, which became defective with the loss of glucose-6-phosphatase as in the case of Osedax. With the emergence of Vestimentifera, TPS was obtained, possibly via transposon-mediated horizontal gene transfer from deep-sea Arthropda, thus gaining the ability of trehalogenensis. The high similarity between the TPS of Vestimentifera and Athropoda suggests that the gene transfer is likely a relatively recent event, which is in line with the phylogenetic position of Vestimentifera as a young member of Siboglinidae.

## Acknowledgements

The authors thank Prof. Ivan Berg (University of Münster, Germany) for valuable suggestions on microbial metabolism. The authors acknowledge the Oceanographic Data Center of IOCAS and the Center for High Performance Computing and System Simulation of Laoshan Laboratory for providing the computing resources. This work was funded by the Strategic Priority Research Program of the Chinese Academy of Sciences (XDA22050000), the National Natural Science Foundation of China (41806202), the Innovation Research Group Project of the National Natural Science Foundation of China (42221005), and the Science Fund Program for Distinguished Young Scholars of Shandong Province (Overseas) (2022HWYQ-087).

## Author contributions

QS, ZY, and LS conceived the study and designed the experiments; QS collected the samples; YS and QS performed the experiments; QS and ZY obtained and analyzed the data; LS, QS, and ZY obtained the funding; QS and ZY wrote the first draft of the manuscript; LS edited the manuscript. All authors have read the manuscript and approved its submission.

## References

1. Cavanaugh, C. M., Gardiner, S. L., Jones, M. L., Jannasch, H. W. & Waterbury, J. B. Prokaryotic cells in the hydrothermal vent tube worm *Riftia Pachyptila* Jones - possible chemoautotrophic symbionts. Science 213, 340–342(1981).

2. Pleijel, F., Dahlgren, T. G. & Rouse, G. W. Progress in systematics: from Siboglinidae to Pogonophora and Vestimentifera and back to Siboglinidae. C R Biol. 332, 140–148(2009).

3. Li Y. et al. Mitogenomics reveals phylogeny and repeated motifs in control regions of the deep-sea family Siboglinidae (Annelida). Mol. Phylogenet. Evol. 85, 221–229(2015).

4. Li Y. et al. Phylogenomics of tubeworms (Siboglinidae, Annelida) and comparative performance of different reconstruction methods. Zool. Scr. 46, 200–213(2017).

5. Rimskaya-Korsakova, N. N., Galkin, S. V. & Malakhov, V. V. The anatomy of the blood vascular system of the giant vestimentiferan tubeworm Riftia pachyptila (Siboglinidae, Annelida). J. Morphol. 278, 810–827(2017).

6. Zimmermann, J. et al. Dual symbiosis with co-occurring sulfur-oxidizing symbionts in vestimentiferan tubeworms from a Mediterranean hydrothermal vent. Environ. Microbiol. 16, 3638–3656(2014).

7. Sun, Y. et al. Genomic signatures supporting the symbiosis and formation of chitinous tube in the deep-sea tubeworm *Paraescarpia echinospica*. Mol. Biol. Evol. 38, 4116–4134(2021).

8. Moggioli, G. et al. Distinct genomic routes underlie transitions to specialised symbiotic lifestyles in deep-sea annelid worms. Nat. Commun. 14, 2814(2023).

9. Wang, M. et al. The genome of a vestimentiferan tubeworm (*Ridgeia piscesae*) provides insights into its adaptation to a deep-sea environment. BMC genomics 24, 72(2023).

10. de Oliveira, A. L., Mitchell. J., Girguis, P. & Bright, M. Novel insights on obligate symbiont lifestyle and adaptation to chemosynthetic environment as revealed by the giant tubeworm genome. Mol. Biol. Evol. 39, msab347(2022).

11. Li Y. et al. Genomic adaptations to chemosymbiosis in the deep-sea seep-dwelling tubeworm Lamellibrachia luymesi. BMC Biol. 17, 91(2019).

12. Lösekann T. et al. Endosymbioses between bacteria and deep-sea siboglinid tubeworms from an Arctic cold seep (Haakon Mosby Mud Volcano, Barents Sea). Environ. Microbiol. 10, 3237–3254 (2008).

13. Hand S. C. Trophosome ultrastructure and the characterization of isolated bacteriocytes from invertebrate-sulfur bacteria symbioses. Biol. Bull. 173, 260–276(1987).

14. Zvi-Kedem, T., Shemesh, E., Tchernov, D. & Rubin-Blum M. The worm affair: fidelity and environmental adaptation in symbiont species that co-occur in vestimentiferan tubeworms. Environ. Microbiol. Rep. 13, 744–52(2021).

15. Yang, Y. et al. Genomic, transcriptomic, and proteomic insights into the symbiosis of deep-sea tubeworm holobionts. ISME J. 14, 135–150(2020).

16. Li Y., Liles M. R. & Halanych K. M. Endosymbiont genomes yield clues of tubeworm success. ISME J. 12, 2785–2795(2018).

17. Reveillaud, J., Anderson, R., Reves-Sohn, S., Cavanaugh, C. & Huber J. A. Metagenomic investigation of vestimentiferan tubeworm endosymbionts from Mid-Cayman Rise reveals new insights into metabolism and diversity. Microbiome 6, 19(2018).

18. Perez, M. & Juniper, K. Insights into symbiont population structure among three vestimentiferan tubeworm host species at eastern Pacific spreading centers. Appl. Environ. Microbiol. 82, 5197–5205(2016).

19. Rubin-Blum, M., Dubilier, N. & Kleiner, M. Genetic evidence for two carbon fixation pathways (the Calvin-Benson-Bassham cycle and the reverse tricarboxylic acid cycle) in symbiotic and free-living bacteria. mSphere 4, e00394–18(2019).

20. Hinzke, T. et al. Host-microbe interactions in the chemosynthetic *Riftia pachyptila* symbiosis. mBio 10, e02243–19(2019).

21. Bright, M. & Sorgo, A. Ultrastructural reinvestigation of the trophosome in adults of *Riftia pachyptila* (Annelida, Siboglinidae). Invertebr. Biol. 122, 345–366(2005).

22. Bright, M., Keckeism H. & Fisher C. R. An autoradiographic examination of carbon fixation, transfer and utilization in the *Riftia pachyptila* symbiosis. Mar. Biol. 136, 621–632(2000).

23. Felbeck, H. & Jarchow, J. Carbon release from purified chemoautotrophic bacterial symbionts of the hydrothermal vent tubeworm *Riftia pachyptila*. Physiol. Zool. 71, 294–302(1998).

24. Sun Q. L., et al. High-throughput sequencing reveals a potentially novel *Sulfurovum* species dominating the microbial communities of the seawater-sediment interface of a deep-sea cold seep in South China Sea. Microorganisms 8, 687(2020).

25. Zhang, J., Sun, Q.-L., Zeng, Z.-G., Chen, S. & Sun, L. Microbial diversity in the deep-sea sediments of Iheya North and Iheya Ridge, Okinawa Trough. Microbiol. Res. 177, 43–52(2015).

26. Bolyen, E. et al. Reproducible, interactive, scalable and extensible microbiome data science using QIIME 2. Nat. Biotechnol. 37, 852–857(2019).

27. Qin, J. et al. A metagenome-wide association study of gut microbiota in type 2 diabetes. Nature 490, 55–60(2012).

28. Fang, C. et al. Assessment of the cPAS-based BGISEQ-500 platform for metagenomic sequencing. Gigascience 7, 1–8(2018).

29. Chen, Y. et al. SOAPnuke: a MapReduce acceleration-supported software for integrated quality control and preprocessing of high-throughput sequencing data. Gigascience 7, 1–6(2018).

30. Luo, R. et al. SOAPdenovo2: an empirically improved memory-efficient short-read de novo assembler. Gigascience 1, 18(2012).

31. Nurk, S., Meleshko, D., Korobeynikov, A., & Pevzner, P.A. metaSPAdes: a new versatile metagenomic assembler. Genome Res. 27, 824–834(2017).

32. Zhu, W., Lomsadze, A. & Borodovsky, M. Ab initio gene identification in metagenomic sequences. Nucleic Acids Res. 38, e132(2010).

33. Fu, L., Niu, B., Zhu, Z., Wu, S. & Li, W. CD-HIT: accelerated for clustering the next-generation sequencing data. Bioinformatics 28, 3150–3152(2012).

34. Uritskiy, G. V., DiRuggiero, J. & Taylor, J. MetaWRAP-a flexible pipeline for genome-resolved metagenomic data analysis. Microbiome 6, 158(2018).

35. Parks, D. H., Imelfort, M., Skennerton, C. T., Hugenholtz, P. & Tyson, G. W. CheckM: assessing the quality of microbial genomes recovered from isolates, single cells, and metagenomes. Genome Res. 25, 1043–1055(2015).

36. Li, D., Liu, C.-M., Luo, R.B., Sadakane, K. & Lam, T.-W. MEGAHIT: an ultra-fast single-node solution for large and complex metagenomics assembly via succinct de Bruijn graph. Bioinformatics. 31,1674–1676(2015).

37. Bray, N.L., Pimentel, H., Melsted, P. & Pachter, L. Near-optimal probabilistic RNA-seq quantification. Nat Biotechnol. 34, 525–527(2016).

38. Laura, V., RAúL, M., Da, S. C. M., & Iraia, F. A new species of lizard of the *L. wiegmannii* group (Iguania: Liolaemidae) from the Uruguayan Savanna. Zootaxa 4294, 443(2017).

39. Heller, P., Casaletto, J., Ruiz, G. & Geller, J. A database of metazoan cytochrome c oxidase subunit I gene sequences derived from GenBank with CO-ARBitrator. Sci Data 5, 1–7(2018).

40. Haiping, L. et al. Draft genome of *Glyptosternon maculatum*, an endemic fish from Tibet Plateau. Gigascience 7, giy104(2018).

41. Liu, B., et al. Estimation of genomic characteristics by analyzing k-mer frequency in de novo genome projects. arXiv 1308.2012(2013). https://arxiv.org/abs/1308.2012.

42. Chin, C.-S., Alexander, D. H., Marks, P., Klammer, A. A. & Korlach, J. Nonhybrid, finished microbial genome assemblies from long-read SMRT sequencing data. Nat. Methods 10, 563(2013).

43. Simão, F., Waterhouse, R. M., Panagiotis, I., Kriventseva, E. V., & Zdobnov, E. M. BUSCO: assessing genome assembly and annotation completeness with single-copy orthologs. Bioinformatics 31, 3210–3212(2015).

44. Durbin, L. R. Fast and accurate short read alignment with Burrows–Wheeler transform. Bioinformatics 25, 1754–1760(2009).

45. Kent, W. J. BLAT—The BLAST-Like Alignment Tool. Genome Res. 12, 656–664(2002).

46. Bao, Z. & Eddy, S. R. Automated de novo identification of repeat sequence families in sequenced genomes. Genome Res. 12, 1269–1276(2002).

47. Price, A. L., Jones, N. C. & Pevzner, P.A. De novo identification of repeat families in large genomes. Bioinformatics **Suppl** 1, i351–8(2005).

48. Bao, W., Kojima, K. K. & Kohany, O. Repbase Update, a database of repetitive elements in eukaryotic genomes. Mob. DNA 6:11(2015).

49. Schffer, A. A., Richa, A., Yu, Y. -K., Michael, G. E. & Altschul, S. F. Composition-based statistics and translated nucleotide searches: Improving the TBLASTN module of BLAST. BMC Biol. 4, 41(2006).

50. Birney, E., Clamp, M. & Durbin, R. GeneWise and Genomewise. Genome Res. 14, 988–995(2004).

51. Mario, S. & Burkhard, M. AUGUSTUS: a web server for gene prediction in eukaryotes that allows user-defined constraints. Nucleic Acids Res. 33, W465–7(2005).

52. Guigó, R., Knudsen, S., Drake, N. & Smith, T. Prediction of gene structure. J. Mol. Biol. 226, 141–157(1992).

53. Burge, C. Prediction of complete gene structures in human genomic DNA. J. Mol. Biol. 268, 78–94(1997).

54. Majoros, W., Pertea, M. & Salzberg, S. TigrScan & GlimmerHMM: two open source ab initio eukaryotic gene-finders. Bioinformatics 20, 2878–2879(2004).

55. Korf, I. Gene finding in novel genomes. BMC bioinformatics 5, 59(2004).

56. Grabherr, M. G. et al. Full-length transcriptome assembly from RNA-Seq data without a reference genome. Nat. Biotechnol. 29, 644–652(2011).

57. Pertea, M., Kim, D., Pertea, G. M., Leek, J. T., & Salzberg, S. L. Transcript-level expression analysis of RNA-seq experiments with HISAT, StringTie and Ballgown. Nat. Protoc. 11, 1650–1667(2016).

58. Kim, D. et al. TopHat2: accurate alignment of transcriptomes in the presence of insertions, deletions and gene fusions. Genome Biol. 14, R36(2013).

59. Pertea, M. et al. StringTie enables improved reconstruction of a transcriptome from RNA-seq reads. Nat. Biotechnol. 33,290–295(2015).

60. Trapnell, C. et al. Differential gene and transcript expression analysis of RNA-seq experiments with TopHat and Cufflinks. Nat. Protoc. 7, 562–578(2012).

61. Haas, B. J. et al. Automated eukaryotic gene structure annotation using EVidenceModeler and the Program to Assemble Spliced Alignments. Genome Biol. 9, R7(2008).

62. Bolger, A. M., Lohse, M., & Usadel, B. Trimmomatic: a flexible trimmer for Illumina sequence data. Bioinformatics 30, 2114–2120(2014).

63. Anders, S., Pyl, P. T., & Huber, W. HTSeq—a Python framework to work with high-throughput sequencing data. Bioinformatics 31,166–169(2015).

64. Ge, S. X., Son, E. W. & Yao, R. iDEP: an integrated web application for differential expression and pathway analysis of RNA-Seq data. BMC bioinformatics 19, 1–24(2018).

65. Sun Q.-L. et al. High temperature-induced proteomic and metabolomic profiles of a thermophilic *Bacillus manusensis* isolated from the deep-sea hydrothermal field of Manus Basin. J. Proteomics 203, 103380(2019).

66. Lutz, R. A. et al. Rapid Growth at Deep-Sea Vents. Nature 371, 663–4(1994).

67. Sorgo, A., Gaill, F., Lechaire, J. P., Arndt, C. & Bright, M. Glycogen storage in the *Riftia pachyptila* trophosome: contribution of host and symbionts. Mar. Ecol. Prog. Ser. 231,115–120(2002).

68. Hinzke, T. et al. Bacterial symbiont subpopulations have different roles in a deep-sea symbiosis. Elife 10, e58371(2021).

69. Shikuma, N.J. Bacteria-stimulated metamorphosis: an ocean of insights from investigating a transient host-microbe interaction. mSystems 6, e00754–21(2021).

70. Miyamoto, T. & Amrein, H. Gluconeogenesis: An ancient biochemical pathway with a new twist. Fly (Austin) 11, 218–223(2017).

71. Matsuda, H., Yamada, T., Yoshida, M. & Nishimura, T. Flies without trehalose. J. Biol. Chem. 290, 1244–1255(2015).

72. Miyamoto, T. & Amrein, H. Neuronal gluconeogenesis regulates systemic glucose homeostasis in *Drosophila melanogaster*. Curr. Biol. 29, 1263–1272 e5(2019).

73. Arguelles, J. C. Thermotolerance and trehalose accumulation induced by heat shock in yeast cells of *Candida albicans*. FEMS Microbiol. Lett. 146, 65–71(1997).

74. Girguis, P. R. & Childress, J. J. Metabolite uptake, stoichiometry and chemoautotrophic function of the hydrothermal vent tubeworm *Riftia pachyptila*: responses to environmental variations in substrate concentrations and temperature. J. Exp. Biol. 209, 3516–3528(2006).

75. Li, Z.-G., Luo, L.-J. & Zhu, L.-P. Involvement of trehalose in hydrogen sulfide donor sodium hydrosulfide-induced the acquisition of heat tolerance in maize (*Zea mays* L.) seedlings. Bot. Stud. 55, 20.

76. Galbraith, J. D., Ludington, A. J., Suh, A., Sanders, K. L. & Adelson, D. L. New environment, new invaders-repeated horizontal transfer of LINEs to sea snakes. Genome Biol. Evol. 12, 2370–2383(2020).

77. Ivancevic, A. M., Kortschak, R. D., Bertozzi, T., & Adelson, D. L. Horizontal transfer of BovB and L1 retrotransposons in eukaryotes. Genome Biol.19, 85 (2018).

78. Sun, Q.-L., Zeng, Z.-G., Chen, S. & Sun, L. First Comparative Analysis of the Community Structures and Carbon Metabolic Pathways of the Bacteria Associated with *Alvinocaris longirostris* in a Hydrothermal Vent of Okinawa Trough. Plos One 11, e0154359(2016).

79. Chen, J.-X., Lyu, Z.-H., Wang, C.-Y., Cheng, J. & Lin, T. RNA interference of a trehalose-6-phosphate synthase gene reveals its roles in the biosynthesis of chitin and lipids in Heortia vitessoides (Lepidoptera: Crambidae). Insect Sci. 27, 212–223(2020).

80. Halanych, K. M., Feldman, R. A. & Vrijenhoek, R. C. Molecular evidence that Sclerolinum brattstromi is closely related to vestimentiferans, not to frenulate pogonophorans (Siboglinidae, Annelida). Biol. Bull. 201, 65–75(2001).

81. Schulze, A. & Halanych, K.M. Siboglinid evolution shaped by habitat preference and sulfide tolerance. Hydrobiologia 496,199–205(2003).

