## Supplementary material for "Integrated multi-approaches reveal unique metabolic mechanisms of Vestimentifera to adapt to deep-sea hydrothermal vents": Figure S1

**Figure S1. The diversity and central carbon metabolism pathways of the *Arcovestia ivanovi* and *Lamellibrachia columna* symbionts.** (**A, B, C**) The sequence tags are classified at the family (A), genus (B), and ASV (C) levels. Each color represents the percentage of the taxon in the total assemblage. A-1 to -9 represent *A. ivanovi* individuals 1 to 9; L-1 to -5 represent *L. columna* individuals 1 to 5. (**D**) Overview of the central carbon metabolisms in *A.* *ivanovi* and *L. columna* symbionts. G, glucose; G-6-P, D-glucose 6-phosphate; F-6-P, D-fructose 6-phosphate; F-1,6-P, D-fructose 1,6-bisphosphate; PGAL, D-glyceraldehyde 3-phosphate; DHAP, glycerone phosphate; 1,3-DPG, 3-phospho-D-glyceroyl phosphate; 3-PGA, 3-phospho-D-glycerate; 2-PGA, 2-phospho-D-glycerate; PEP, phosphoenolpyruvate; Pyru, pyruvate; Ace-CoA, acetyl-CoA; Rbu-1,5-P, D-ribulose 1,5-bisphosphate; Rbu-5-P, D-ribulose 5-phosphate; Rbo-5-P, D-ribose 5-phosphate; Sedo-7P, sedoheptulose 7-phosphate; Sedo-1,7-P, sedoheptulose 1,7-bisphosphate; Ery-4P, D-erythrose 4-phosphate; G-1-P, D-glucose 1-phosphate; ADP-G, ADP-glucose; Amy, amylose; Glyco, glycongen; Oxalo, oxaloacetate; Mal, malate; Fum, fumarate; Suc, succinate; Suc-CoA, succinyl-CoA; 2-oxog, 2-oxoglutarate; Isocit, isocitrate; Aco, aconitate; Cit, citrate; 1, ribulose-bisphosphate carboxylase; 2, phosphoglycerate kinase; 3, glyceraldehyde 3-phosphate dehydrogenase ; 4, fructose-bisphosphate aldolase; 5, ATP-dependent phosphofructokinase; 6, transketolase; 7, triosephosphate isomerase; 8, fructose-bisphosphate aldolase; 9, ATP-dependent phosphofructokinase; 10, transketolase; 11, ribose 5-phosphate isomerase A; 12, ribulose-phosphate 3-epimerase; 13, phosphoribulokinase; 14, 2,3-bisphosphoglycerate-independent phosphoglycerate mutase; 15, enolase; 16, pyruvate, water dikinase; 17, pyruvate ferredoxin oxidoreductase; 18, citrate synthase; 19, malate dehydrogenase; 20, fumarate hydratase; 21, succinate dehydrogenase / fumarate reductase; 22, succinate-CoA ligase; 23, 2-oxoglutarate ferredoxin oxidoreductase; 24, isocitrate dehydrogenase; 25/26, aconitate hydratase; 27, citrate lyase; 28, glucose-6-phosphate isomerase; 29, glucokinase; 30, phosphoglucomutase; 31, glucose-1-phosphate adenylyltransferase; 32, starch synthase; 33, 1,4-alpha-glucan branching enzyme; 34, pyruvate carboxylase. The solid and dashed lines represent the reactions that can and cannot, respectively, occur in vestimentiferan symbionts. (**E**) The glycogen pathway in siboglinid symbionts. GlgC, glucose-1-phosphate adenylyltransferase; GlgA, starch synthase; GlgB, 1,4-alpha-glucan branching enzyme. +, present; -, absent.

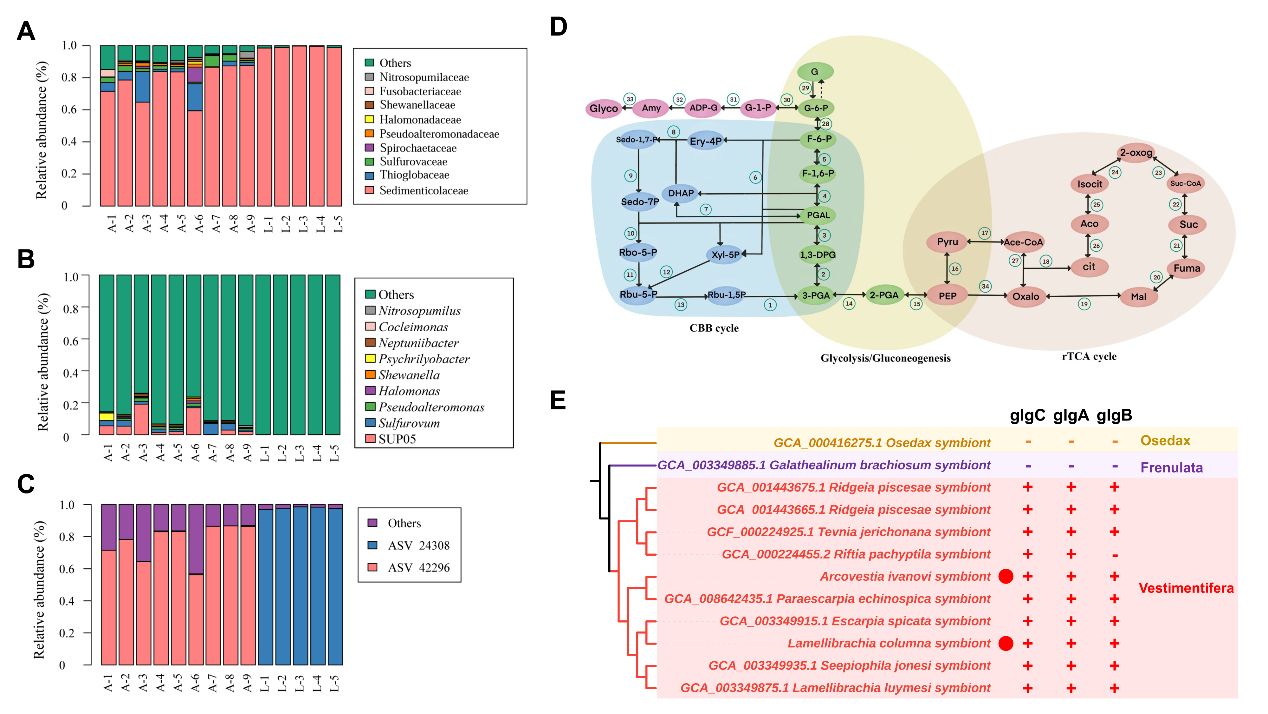
