## Supplementary material for "Integrated multi-approaches reveal unique metabolic mechanisms of Vestimentifera to adapt to deep-sea hydrothermal vents": Figure S2

**Figure S2. The expression of the trehalose 6-phosphate synthase gene in vestimentiferans.** Blue, red, and green represent plume, vestimentum, and trophosome, respectively. The Y-axis represents the expression CPM at Ln scale.

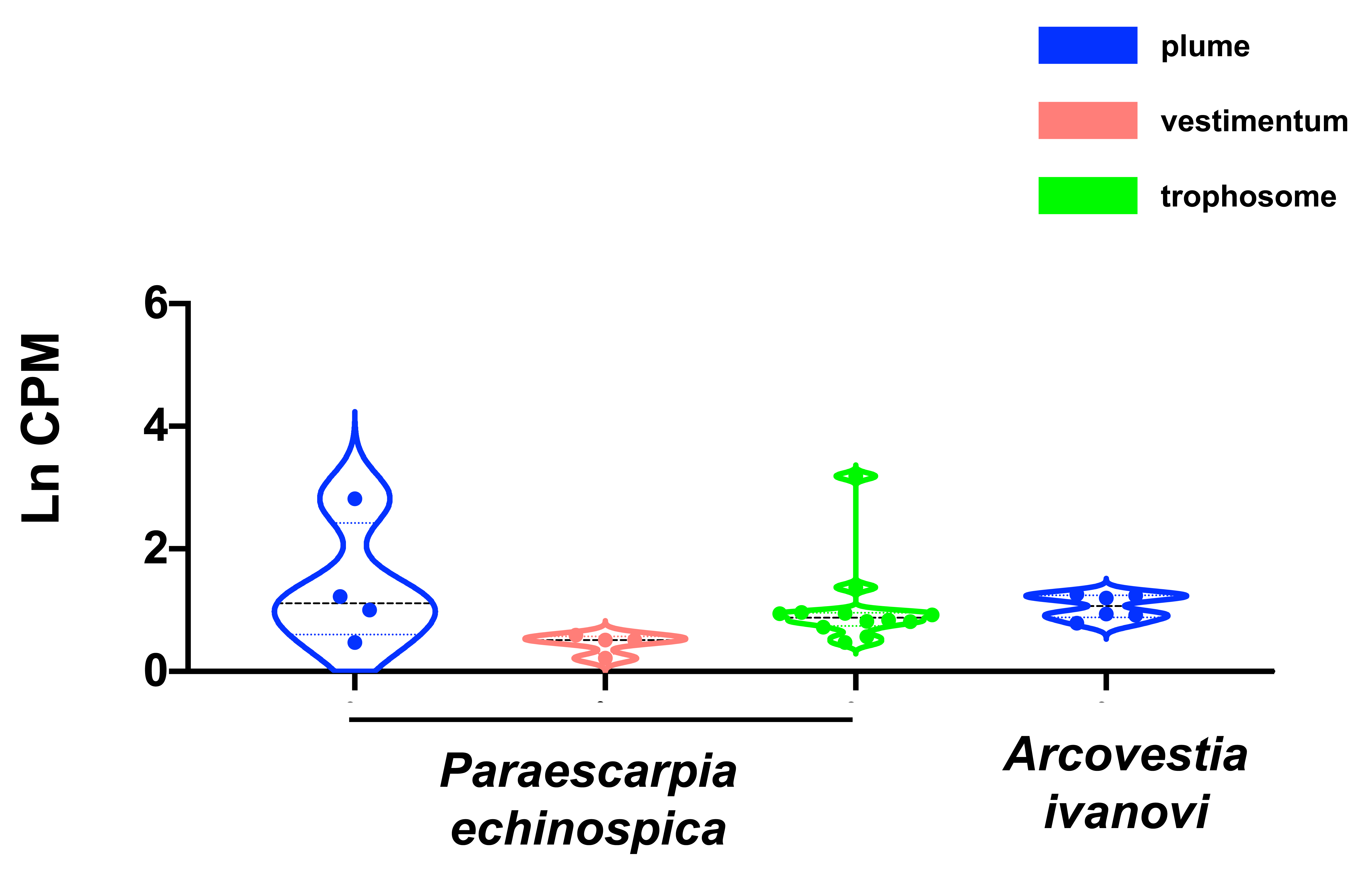
