## Supplementary material for "Integrated multi-approaches reveal unique metabolic mechanisms of Vestimentifera to adapt to deep-sea hydrothermal vents": Table S1

**Table S1. Expression analysis of the central carbon metabolism genes in the symbionts of *Arcovestia ivanovi* (A1 and A2) and *Lamellibrachia columna* (L1).** Transcripts per million (TPM) was used to assess the gene expression level.

| No. | KEGG orthology | Enzyme | Gene | A1 | A2 | L1 |
| --- | --- | --- | --- | --- | --- | --- |
| **The CBB cycle** | | | | | | |
| 1 | K01601 | ribulose-bisphosphate carboxylase large chain | *cbbL* | 116.26 | 81.22 | 1986.90 |
| 2 | K00927 | phosphoglycerate kinase | *pgk* | 33.14 | 19.85 | 2.41 |
| 3 | K00134 | glyceraldehyde 3-phosphate dehydrogenase | *gapdh* | 356.00 | 212.81 | 32.61 |
| 4 | K01623 | fructose-bisphosphate aldolase | *fba* | 175.10 | 73.02 | 21.01 |
| 5 | K21071 | ATP-dependent phosphofructokinase | *pfk* | 53.61 | 15.91 | 45.33 |
| 6 | K00615 | transketolase | *tkt* | 56.93 | 32.27 | 5.85 |
| 7 | K01803 | triosephosphate isomerase | *tpi* | 16.01 | 3.24 | 17.79 |
| 8 | K01624 | fructose-bisphosphate aldolase | *fba* | 175.10 | 73.02 | 21.01 |
| 9 | K21071 | ATP-dependent phosphofructokinase | *pfk* | 53.61 | 15.91 | 45.33 |
| 10 | K00615 | transketolase | *tkt* | 56.93 | 32.27 | 5.85 |
| 11 | K01807 | ribose 5-phosphate isomerase | *rpiA* | 21.41 | 4.09 | 10.41 |
| 12 | K01783 | ribulose-phosphate 3-epimerase | *rpe* | 139.44 | 32.05 | 36.63 |
| 13 | K00855 | phosphoribulokinase | *prk* | 68.65 | 47.46 | 62.09 |
| 28 | K01810 | glucose-6-phosphate isomerase | *gpi* | 1.19 | 0.62 | 43.85 |
| **Gycolysis/glyconeogenesis** | | | | | | |
| 29 | K00845 | glucokinase | *glk* | 0.68 | 0.30 | 4.21 |
| 28 | K01810 | glucose-6-phosphate isomerase | *gpi* | 1.19 | 0.62 | 43.85 |
| 5 | K21071 | diphosphate-dependent phosphofructokinase | *pfk* | 53.61 | 15.91 | 45.33 |
| 4 | K01624 | fructose-bisphosphate aldolase | *fba* | 175.10 | 73.02 | 21.01 |
| 3 | k00134 | glyceraldehyde 3-phosphate dehydrogenase | *gapdh* | 356.00 | 212.81 | 32.61 |
| 7 | K01803 | triosephosphate isomerase | *tpi* | 16.01 | 3.24 | 17.79 |
| 2 | K00927 | phosphoglycerate kinase | *pgk* | 33.14 | 19.85 | 2.41 |
| 14 | K15633 | 2,3-bisphosphoglycerate-independent phosphoglycerate mutase | *gpmI* | 4.22 | 1.48 | 8.04 |
| 15 | K01689 | enolase | *eno* | 6.98 | 1.92 | 3.62 |
| 16 | K00873 | pyruvate kinase | *pk* | 13.73 | 7.59 | 3.53 |
| **The TCA/rTCA cycle** | | | | | | |
| 17 | K03737 | pyruvate ferredoxin oxidoreductase | *por* | 2.37 | 3.26 | 3.03 |
| 16 | K01006 | pyruvate, water dikinase | *ppsA* | 6.82 | 3.68 | 3.46 |
| 34 | K01959 | pyruvate carboxylase subunit A | *pycA* | 10.78 | 2.50 | 8.86 |
|  | K01960 | pyruvate carboxylase subunit B | *pycB* | 8.02 | 3.35 | 11.68 |
| 19 | K00024 | malate dehydrogenase | *mdh* | 20.92 | 11.40 | 22.67 |
| 20 | K01676 | fumarate hydratase | *fum* | 4.27 | 1.62 | 8.68 |
| 21 | K00239 | succinate dehydrogenase / fumarate reductase flavoprotein subunit | *frdA* | 5.74 | 8.74 | 49.20 |
|  | K00240 | succinate dehydrogenase / fumarate reductase iron-sulfur subunit | *frdB* | 8.72 | 3.46 | 116.21 |
| 22 | K01902 | succinate--CoA ligase subunit alpha | *sucD* | 11.19 | 5.66 | 94.69 |
|  | K01903 | ADP-forming succinate--CoA ligase subunit beta | *sucC* | 5.05 | 3.16 | 33.65 |
| 23 | K00174 | 2-oxoglutarate ferredoxin oxidoreductase subunit alpha | *korA* | 7.30 | 4.19 | 47.75 |
|  | K00175 | 2-oxoglutarate oxidoreductase | *korB* | 19.28 | 14.24 | 130.72 |
| 24 | K00031 | isocitrate dehydrogenase | *icd* | 19.25 | 11.88 | 37.90 |
| 25/26 | K01681 | aconitate hydratase | *aco* | 23.57 | 8.87 | 41.34 |
| 27 | K15231 | ATP-dependent citrate lyase | *aclAB* | 18.00 | 14.06 | 25.28 |
| 18 | K01647 | citrate synthase | *cs* | 1.27 | 0.80 | 3.65 |
| **Glycogen biosynthesis** | | | | | | |
| 30 | K01835 | phosphoglucomutase | *pgm* | 6.24 | 2.11 | 3.27 |
| 31 | K00975 | glucose-1-phosphate adenylyltransferase | *glgC* | 20.30 | 15.51 | 26.28 |
| 32 | K00703 | starch synthase | *glgA* | 2.61 | 1.11 | 1.69 |
| 33 | K00700 | 1,4-alpha-glucan branching enzyme | *glgB* | 11.61 | 5.93 | 8.34 |
| **Riboflavin biosynthesis** | | | | | | |
| 34 | K02858 | 3,4-dihydroxy 2-butanone 4-phosphate synthase | *ribB* | 2.98 | 0.78 | 7.05 |
| 35 | K00794 | 6,7-dimethyl-8-ribityllumazine synthase | *ribH* | 10.14 | 6.83 | 10.73 |
| 36 | K00793 | riboflavin synthase | *ribE* | 5.90 | 3.65 | 9.85 |
