## Supplementary material for "Integrated multi-approaches reveal unique metabolic mechanisms of Vestimentifera to adapt to deep-sea hydrothermal vents": Table S2

**Table S2. The metabolites detected in the metabolomics of *Lamellibrachia columna*.** Fold change indicates the ratio of the metabolites in the trophosome and vestimentitum of the tubeworm. Statistical significance was determined with unpaired Student’s t test.

| No. | **name** | **adduct** | **description** | **fold change** | **p-value** |
| --- | --- | --- | --- | --- | --- |
| 1 | M377T211_2 | (M+H)+ | Riboflavin | 1247.251679 | 0.000919 |
| 2 | M241T159 | (M-H)- | Lumichrome | 203.6112617 | 0.00856 |
| 3 | M173T342 | (M+H)+ | Glycerol 3-phosphate | 93.77069496 | 0.011201 |
| 4 | M241T97 | (M-H)- | Thymidine | 73.65079456 | 0.087985 |
| 5 | M190T186 | (M+H)+ | Kynurenic acid | 54.51155391 | 0.033612 |
| 6 | M314T397 | M- | Geranyl diphosphate | 49.23835137 | 0.151591 |
| 7 | M401T395 | (M+CH3COO)- | Galactinol | 44.00156896 | 0.112474 |
| 8 | M283T307 | (M-H)- | Xanthosine | 37.13333514 | 0.017672 |
| 9 | M252T140 | (M+H)+ | Deoxyadenosine | 36.68946874 | 0.266626 |
| 10 | M153T308 | (M+H)+ | Xanthine | 35.4525489 | 0.02508 |
| 11 | M146T37_1 | (M+H-2H2O)+ | DL-O-tyrosine | 31.13021773 | 0.009845 |
| 12 | M182T75 | (M-H)- | L-homocysteic acid | 28.37162567 | 0.000772 |
| 13 | M172T38 | (M-H2O-H)- | 5-Hydroxyindoleacetate | 28.30043033 | 0.014588 |
| 14 | M219T276_2 | (M-H)- | 5-Hydroxytryptophan | 27.08049372 | 0.026313 |
| 15 | M298T194 | (M+H)+ | 2-Methylguanosine | 25.00656445 | 0.071766 |
| 16 | M221T276_1 | (M+NH4)+ | Indole-3-pyruvic acid | 23.5001045 | 0.034179 |
| 17 | M200T453 | (M+H)+ | O-Phospho-L-homoserine | 22.5199193 | 0.02376 |
| 18 | M202T35 | (M-H)- | 3-(3-Indolyl)-2-oxopropanoic acid | 21.32295273 | 0.000132 |
| 19 | M158T47 | (M+H-H2O)+ | Indoleacetic acid | 20.09014714 | 0.000252 |
| 20 | M311T156 | (M+H)+ | Methoprene (S) | 19.7215088 | 0.001036 |
| 21 | M266T236 | (M-2H+3K)+ | (-)-Norephedrine | 19.13995081 | 0.029776 |
| 22 | M341T394 | (M-H)- | Trehalose | 18.96148259 | 0.009601 |
| 23 | M157T46 | (M+H)+ | 3-Indoleacetonitrile | 17.73745343 | 0.000823 |
| 24 | M138T282_2 | M+ | Trigonelline | 16.79435485 | 0.015011 |
| 25 | M120T435 | (M+H-H2O)+ | Anthranilic acid (Vitamin L1) | 16.69738642 | 0.050666 |
| 26 | M253T43 | (M-H)- | cis-9-Palmitoleic acid | 15.08534213 | 0.038027 |
| 27 | M121T435 | (M+H-H2O)+ | Gentisaldehyde | 14.53734295 | 0.000372 |
| 28 | M144T384 | (M+H-H2O)+ | Indole-3-carboxylic acid | 11.52083083 | 0.007522 |
| 29 | M282T129 | (M+H)+ | N6-methyladenosine | 11.31152204 | 0.028293 |
| 30 | M142T28 | (M+Na-2H)- | L-Cysteine | 10.83617434 | 9.56E-05 |
| 31 | M175T306 | (M+CH3COO)- | alpha-ketoisovaleric acid | 10.74332426 | 0.000702 |
| 32 | M296T105 | M+ | Tyr-Asp | 10.66511857 | 0.120288 |
| 33 | M366T168 | (M+NH4)+ | Camptothecin | 9.876660968 | 5.85E-05 |
| 34 | M113T160 | (M+H)+ | Uracil | 9.04202824 | 0.005937 |
| 35 | M341T189 | (M+H-H2O)+ | 1-Stearoyl-sn-glycerol | 8.646618702 | 0.305311 |
| 36 | M213T342 | (M-H)- | 1-Deoxy-D-xylulose 5-phosphate | 7.811952314 | 0.014376 |
| 37 | M251T178 | (M-H)- | Deoxyinosine | 7.685804482 | 0.01505 |
| 38 | M326T152 | (M-H+2Na)+ | 2'-O-methyladenosine | 7.166690473 | 0.016672 |
| 39 | M295T342 | (M+H)+ | gamma-L-Glutamyl-L-phenylalanine | 7.069504079 | 0.009853 |
| 40 | M324T454 | (M-2H+3Na)+ | 2'-O-methylcytidine | 6.973756898 | 0.005371 |
| 41 | M333T442_2 | (M+H)+ | Lys-Trp | 6.972718817 | 0.017664 |
| 42 | M409T459 | (M-H+2Na)+ | Xanthylic acid (XMP) | 6.884500785 | 5.03E-05 |
| 43 | M359T98 | (M+H-H2O)+ | (-)-Riboflavin | 6.838679486 | 0.013835 |
| 44 | M510T191 | (M+H)+ | 1-O-Octadecyl-sn-glyceryl-3-phosphorylcholine | 6.74593869 | 0.102475 |
| 45 | M178T245 | (M+K-2H)- | Ethosuximide | 6.658103927 | 0.001174 |
| 46 | M302T121 | (M+H)+ | Sphinganine | 6.63278543 | 0.039139 |
| 47 | M137T178 | (M+H)+ | Hypoxanthine | 6.183130757 | 0.042694 |
| 48 | M336T460 | (M+H)+ | Nicotinate D-ribonucleotide | 6.089489945 | 0.005074 |
| 49 | M266T228 | (M-H)- | Deoxyguanosine | 5.739578222 | 0.034506 |
| 50 | M550T184 | M+ | 1-O-(cis-9-Octadecenyl)-2-O-acetyl-sn-glycero-3-phosphocholine | 5.692034785 | 0.086498 |
| 51 | M89T299 | (M-H)- | DL-lactate | 5.671404207 | 0.002318 |
| 52 | M164T55 | (M-H)- | Formylanthranilic acid | 5.640403997 | 0.284497 |
| 53 | M306T438 | (M-H)- | 2'-Deoxycytidine 5'-monophosphate (dCMP) | 5.451616146 | 0.083657 |
| 54 | M236T253 | (M-H)- | Biopterin | 5.303316588 | 0.011492 |
| 55 | M152T227_2 | (M+H)+ | 2-Hydroxyadenine | 4.863227909 | 0.020589 |
| 56 | M216T387 | (M+H)+ | sn-Glycerol 3-phosphoethanolamine | 4.647655285 | 0.040612 |
| 57 | M197T192 | (M+Na-2H)- | L-Ascorbic acid | 4.514842558 | 0.001169 |
| 58 | M546T186_2 | (M+Na)+ | 1-Stearoyl-sn-glycerol 3-phosphocholine | 4.496201158 | 0.092535 |
| 59 | M168T95 | (M+H)+ | Pyridoxal (Vitamin B6) | 4.3803069 | 1.09E-05 |
| 60 | M330T266 | (M+H)+ | Adenosine 2',3'-cyclic monophosphate | 4.234550811 | 8.53E-05 |
| 61 | M563T446 | (M+CH3COO)- | Raffinose | 4.026565206 | 0.005456 |
| 62 | M170T38 | (M+H)+ | Pyridoxine | 4.011638252 | 0.020716 |
| 63 | M455T390_1 | (M-H)- | Flavin mononucleotide (FMN) | 4.010002454 | 0.015001 |
| 64 | M218T269 | (M-H)- | Pantothenate | 4.009854971 | 1.47E-08 |
| 65 | M168T361 | (M-H)- | L-Cysteic acid | 3.99972463 | 0.103608 |
| 66 | M318T36 | (M-H)- | Chloroquine | 3.879693613 | 0.125458 |
| 67 | M175T531 | (M+H)+ | L-Arginine | 3.810165544 | 1.47E-05 |
| 68 | M260T529 | (M+H)+ | Lys-Leu | 3.804117066 | 0.130962 |
| 69 | M132T397_2 | (M-H)- | D-Aspartic acid | 3.7847742 | 1.53E-05 |
| 70 | M62T387 | (M+H)+ | Ethanolamine | 3.656000139 | 0.047439 |
| 71 | M786T393 | (M+H)+ | Flavin adenine dinucleotide (FAD) | 3.51516375 | 0.000322 |
| 72 | M112T438 | (M+H)+ | Cytosine | 3.447792218 | 0.066155 |
| 73 | M827T138_1 | M+ | PC(20:5(5Z,8Z,11Z,14Z,17Z)/20:5(5Z,8Z,11Z,14Z,17Z)) | 3.34619273 | 0.231213 |
| 74 | M158T508 | (M+H-H2O)+ | L-Citrulline | 3.302227019 | 0.013773 |
| 75 | M130T64 | (M+H)+ | .beta.-Homoproline | 3.171734081 | 0.049799 |
| 76 | M161T530 | (M+H)+ | N6-Methyl-L-lysine | 3.105387056 | 0.003487 |
| 77 | M188T383 | (M-H)- | N-Acetyl-L-glutamate | 3.020452061 | 0.000494 |
| 78 | M345T408 | (M+Na)+ | Thymidine 5'-monophosphate | 3.016383514 | 0.080737 |
| 79 | M332T414 | (M+H)+ | 2'-Deoxyadenosine 5'-monophosphate (dAMP) | 2.94537613 | 0.093677 |
| 80 | M175T294 | (M+H)+ | DL-Arginine | 2.88205817 | 0.221572 |
| 81 | M219T299 | (M+NH4)+ | Cysteine-S-sulfate | 2.832344967 | 6.42E-05 |
| 82 | M462T189 | (M+H)+ | Psychosine | 2.831909544 | 0.227459 |
| 83 | M123T60 | (M+H)+ | Nicotinamide | 2.822690619 | 0.016565 |
| 84 | M134T398 | (M+H)+ | L-Aspartate | 2.805252495 | 0.001188 |
| 85 | M185T328 | (M+CH3COO)- | Thymine | 2.751344751 | 0.083328 |
| 86 | M522T189_3 | (M+H)+ | 1-Oleoyl-sn-glycero-3-phosphocholine | 2.702513878 | 0.152333 |
| 87 | M611T409 | (M-H)- | Novobiocin | 2.666776764 | 0.006387 |
| 88 | M175T338 | (M-H)- | 2-Isopropylmalic acid | 2.633660388 | 3.63E-05 |
| 89 | M275T572 | (M+H)+ | Lys-Lys | 2.566563745 | 0.014524 |
| 90 | M173T239 | (M+CH3CN+H)+ | L-Isoleucine | 2.494054905 | 0.141429 |
| 91 | M327T40 | (M-H)- | (4Z,7Z,10Z,13Z,16Z,19Z)-4,7,10,13,1 6,19-Docosahexaenoic acid | 2.449284504 | 0.311299 |
| 92 | M321T430 | (M-H)- | Deoxythymidine 5'-phosphate (dTMP) | 2.428102999 | 0.116535 |
| 93 | M295T42 | (M-H)- | 13(S)-HODE | 2.417755862 | 0.093225 |
| 94 | M348T404 | (M+H)+ | Adenosine 3'-monophosphate | 2.408035534 | 0.001783 |
| 95 | M322T431 | (M-H)- | Cytidine 5'-monophosphate (CMP) | 2.308440642 | 0.025324 |
| 96 | M323T437 | (M-H)- | Uridine 5'-monophosphate (UMP) | 2.265188543 | 0.062938 |
| 97 | M244T442 | (M+CH3COO)- | 3-Phosphoserine | 2.192624012 | 0.438854 |
| 98 | M274T377_2 | (M+H)+ | Arg-Val | 2.182601783 | 0.133113 |
| 99 | M318T415 | M- | Lecanoric acid | 2.176722949 | 0.012587 |
| 100 | M254T247 | (M+H)+ | D-Neopterin | 2.15195586 | 0.33306 |
| 101 | M198T185 | (M+CH3CN+H)+ | Orotate | 2.136061602 | 0.274264 |
| 102 | M206T478 | (M+Na)+ | Phosphorylcholine | 2.119127558 | 0.004705 |
| 103 | M294T170_1 | (M+CH3CN+H)+ | Tyr-Ala | 2.073241831 | 0.339064 |
| 104 | M496T191_3 | (M+H)+ | 1-Palmitoyl-sn-glycero-3-phosphocholine | 2.023785322 | 0.232219 |
| 105 | M506T195 | (M+H-H2O)+ | 1-Stearoyl-2-hydroxy-sn-glycero-3-phosphocholine | 2.013215713 | 0.193777 |
| 106 | M188T255_2 | (M+H-H2O)+ | DL-Indole-3-lactic acid | 2.003670475 | 0.00858 |
| 107 | M359T387 | (2M-H)- | D-(+)-Mannose | 1.986199452 | 0.009324 |
| 108 | M191T469 | (M+H)+ | Diaminopimelic acid | 1.980245559 | 0.280048 |
| 109 | M290T401 | (M+CH3COO+2H)+ | Pro-Asn | 1.972201751 | 0.211415 |
| 110 | M301T32 | (M+H-H2O)+ | 12-oxo-ETE | 1.92999909 | 0.036657 |
| 111 | M128T217 | (M+H-H2O)+ | 4-Guanidinobutyric acid | 1.866461387 | 0.354718 |
| 112 | M228T178 | (M+H)+ | Deoxycytidine | 1.853582048 | 0.012695 |
| 113 | M303T565 | (M+H)+ | Lys-Arg | 1.831135151 | 0.122525 |
| 114 | M191T333 | (M-H)- | Quinate | 1.819178575 | 0.00505 |
| 115 | M290T450 | (M+H)+ | Arg-Asp | 1.805812038 | 0.0883 |
| 116 | M205T454 | (M-H)- | Homocitrate | 1.803814015 | 0.213901 |
| 117 | M141T348 | (M-H2O-H)- | 2-Oxoadipic acid | 1.792675293 | 0.00119 |
| 118 | M116T255 | (M-H)- | Indole | 1.782797587 | 0.011768 |
| 119 | M403T461 | (M+NH4-2H)- | 2'-Deoxycytidine diphosphate (dCDP) | 1.756788419 | 0.051385 |
| 120 | M154T458 | (M+NH4-2H)- | Fosfomycin | 1.754267293 | 0.007485 |
| 121 | M145T367_2 | (M-H)- | L-Glutamine | 1.746840532 | 0.002016 |
| 122 | M472T166 | (M-H+2Na)+ | Stearoylcarnitine | 1.744519784 | 0.449686 |
| 123 | M203T255 | (M-H)- | L-Tryptophan | 1.725951045 | 0.04066 |
| 124 | M164T359_2 | (M-H)- | DL-Methionine sulfoxide | 1.707062485 | 0.214882 |
| 125 | M313T250_2 | (M+H-H2O)+ | 1-Palmitoylglycerol | 1.699823322 | 0.102098 |
| 126 | M135T103 | (M-H)- | 2-Methylbenzoic acid | 1.695712753 | 0.02541 |
| 127 | M127T454 | (M-H)- | Dihydrothymine | 1.687599739 | 0.004919 |
| 128 | M269T214 | (M+H)+ | Allopurinol riboside | 1.634546568 | 0.23019 |
| 129 | M142T337 | (M+H)+ | L-Histidinol | 1.629024271 | 0.315185 |
| 130 | M298T94 | (M+H)+ | S-Methyl-5'-thioadenosine | 1.612884445 | 0.145133 |
| 131 | M177T303 | (M-H)- | 2-Dehydro-3-deoxy-D-gluconate | 1.571065131 | 0.201853 |
| 132 | M152T103 | (M+H)+ | DL-.alpha.-Phenylglycine | 1.509110436 | 0.149529 |
| 133 | M468T194 | (M+H)+ | 1-Myristoyl-sn-glycero-3-phosphocholine | 1.487545812 | 0.46718 |
| 134 | M272T443 | (M+H)+ | Pro-Arg | 1.481475235 | 0.318593 |
| 135 | M227T45 | (M-H)- | Myristic acid | 1.480810664 | 0.381903 |
| 136 | M258T339_2 | (M-H)- | D-Glucosamine 1-phosphate (Glucosamine-1P) | 1.451756905 | 0.238662 |
| 137 | M120T253_2 | (M+H-H2O)+ | Tyramine | 1.438507103 | 0.093083 |
| 138 | M149T253 | (M+H-H2O)+ | Phenyllactic acid | 1.42660075 | 0.113162 |
| 139 | M166T253_2 | (M+H)+ | L-Phenylalanine | 1.41894876 | 0.098733 |
| 140 | M244T364 | M- | L-Fucose-1-phosphate | 1.399674946 | 0.229324 |
| 141 | M294T37 | (M+CH3CN+H)+ | Ala-Tyr | 1.376969787 | 0.195884 |
| 142 | M124T217 | (M+H)+ | Nicotinate | 1.365192004 | 0.29027 |
| 143 | M294T35 | (M+NH4)+ | Stearidonic Acid | 1.356623662 | 0.35569 |
| 144 | M339T184 | (M+H)+ | (+-)8,9-DHET | 1.354450636 | 0.623575 |
| 145 | M522T380 | (M+NH4)+ | Maltotriose | 1.343971534 | 0.342863 |
| 146 | M329T39 | (M-H)- | 7Z, 10Z, 13Z, 16Z, 19Z-Docosapentaenoic acid | 1.339789101 | 0.466638 |
| 147 | M235T138 | (M+H-H2O)+ | His-Pro | 1.336013853 | 0.432358 |
| 148 | M130T258 | (M-H)- | L-Leucine | 1.311211605 | 0.002943 |
| 149 | M204T449 | (M+H)+ | Gly-Lys | 1.307114017 | 0.238721 |
| 150 | M267T215 | (M-H)- | Inosine | 1.303779177 | 0.595841 |
| 151 | M304T445_2 | (M+H)+ | Arg-Glu | 1.282091738 | 0.417127 |
| 152 | M227T350 | (2M+K)+ | Dimethyl sulfone | 1.280704871 | 0.125211 |
| 153 | M182T296_2 | (M+H)+ | L-Tyrosine | 1.264945082 | 0.211249 |
| 154 | M279T41 | (M-H)- | Linoleic acid | 1.255383688 | 0.638552 |
| 155 | M195T376 | (M-H)- | Galactonic acid | 1.243393875 | 0.569067 |
| 156 | M246T390 | (M+H)+ | Lys-Val | 1.237541313 | 0.518326 |
| 157 | M95T296 | (M+H)+ | Phenol | 1.236075889 | 0.196159 |
| 158 | M102T392_2 | (M-H)- | (S)-2-aminobutyric acid | 1.189328959 | 0.231059 |
| 159 | M119T168 | (M+H-H2O)+ | L-Threonate | 1.17937244 | 0.430158 |
| 160 | M277T42_2 | (M-H)- | all cis-(6,9,12)-Linolenic acid | 1.178232422 | 0.686445 |
| 161 | M230T380_1 | (M-H2O-H)- | Pyridoxine 5-phosphate | 1.168924651 | 0.698196 |
| 162 | M341T369 | (M-H)- | D-Maltose | 1.167599483 | 0.601958 |
| 163 | M258T469 | M+ | Glycerophosphocholine | 1.166203149 | 0.754707 |
| 164 | M296T35_2 | (M+NH4)+ | alpha-Linolenic acid | 1.156810577 | 0.584192 |
| 165 | M228T392 | (M+CH3CN+Na)+ | 2-Phenylbutyric acid | 1.153234914 | 0.169831 |
| 166 | M213T429 | (M+H-H2O)+ | Pro-Asp | 1.125830857 | 0.386899 |
| 167 | M268T168_2 | (M+H)+ | Adenosine | 1.123105088 | 0.638898 |
| 168 | M133T113 | (M-H2O-H)- | Ribitol | 1.117608112 | 0.790698 |
| 169 | M355T231 | (M-H)- | Estradiol valerate | 1.106272045 | 0.540173 |
| 170 | M147T296 | (M+H-H2O)+ | 4-Hydroxycinnamic acid | 1.104882962 | 0.598264 |
| 171 | M383T376 | (M-H)- | S-Adenosyl-L-homocysteine | 1.09911017 | 0.532802 |
| 172 | M145T517 | (M-H)- | L-Lysine | 1.095388089 | 0.741282 |
| 173 | M174T390 | (M-H)- | N-Acetyl-L-aspartic acid | 1.094598344 | 0.710957 |
| 174 | M222T255 | (M+H)+ | N-Acetylmannosamine | 1.085472817 | 0.800032 |
| 175 | M684T442 | (M+NH4)+ | Glycogen | 1.079769532 | 0.779905 |
| 176 | M136T296_2 | (M+H-H2O)+ | Dopamine | 1.071468242 | 0.759974 |
| 177 | M133T370 | (M+H)+ | L-Asparagine | 1.068034646 | 0.813778 |
| 178 | M303T443 | (M+CH3COO)- | Uridine | 1.062372495 | 0.878844 |
| 179 | M129T53 | (M-H)- | ketoisocaproic acid | 1.057037862 | 0.880477 |
| 180 | M312T453 | (M+H)+ | His-Arg | 1.04536408 | 0.924918 |
| 181 | M118T268_2 | (M+H)+ | Betaine | 1.035486211 | 0.828876 |
| 182 | M252T111 | (M+H)+ | 5'-Deoxyadenosine | 1.018945107 | 0.966067 |
| 183 | M103T387 | (M-H)- | Malonic acid | 1.006071131 | 0.971701 |
| 184 | M114T43 | (M+H)+ | epsilon-Caprolactam | 0.991625554 | 0.932868 |
| 185 | M312T168 | M+ | Tyr-Met | 0.984694342 | 0.933537 |
| 186 | M231T328 | (M+H-H2O)+ | Thr-Glu | 0.983634327 | 0.944147 |
| 187 | M285T408 | (M+H)+ | Glu-His | 0.980856752 | 0.951578 |
| 188 | M194T334 | (M+CH3CN+H)+ | 2-Acetylresorcinol | 0.976128259 | 0.973123 |
| 189 | M215T330 | (M+H)+ | Pro-Val | 0.973480272 | 0.915146 |
| 190 | M391T150 | (M-H)- | Chenodeoxycholate | 0.971277347 | 0.961218 |
| 191 | M128T298 | (M+H-H2O)+ | 4-acetamidobutanoate | 0.962156776 | 0.90182 |
| 192 | M542T429 | (M+H-H2O)+ | ADP-ribose | 0.956762805 | 0.839697 |
| 193 | M210T253 | (M+CH3CN+Na)+ | Coumarin | 0.950252211 | 0.656888 |
| 194 | M263T223 | (M-H)- | Alpha-N-Phenylacetyl-L-glutamine | 0.940484133 | 0.900169 |
| 195 | M159T400 | (M+K-2H)- | D-Threitol | 0.934847361 | 0.409117 |
| 196 | M301T39 | (M-H)- | Eicosapentaenoic Acid | 0.930722676 | 0.883208 |
| 197 | M176T189 | (M-H)- | N-Formylmethionine | 0.913456174 | 0.767164 |
| 198 | M117T383_2 | (M-H)- | Succinate | 0.912833648 | 0.414967 |
| 199 | M73T383_2 | (M-H)- | Propionic acid | 0.912411462 | 0.418261 |
| 200 | M399T467 | (M+H)+ | S-Adenosylmethionine | 0.897302029 | 0.703484 |
| 201 | M114T370_1 | (M-H)- | Maleamic acid | 0.879160627 | 0.598713 |
| 202 | M244T455 | (M+CH3CN+H)+ | Pro-Ser | 0.8624496 | 0.664654 |
| 203 | M131T360_2 | (M-H)- | Glutaric acid | 0.822447785 | 0.490939 |
| 204 | M235T424 | (M+H)+ | Glu-Ser | 0.812252184 | 0.170444 |
| 205 | M189T499 | (M+H)+ | L-NG-Monomethylarginine | 0.811328543 | 0.596964 |
| 206 | M275T447 | (M+CH3CN+H)+ | Lys-Ser | 0.810063499 | 0.528643 |
| 207 | M489T437 | (M+H)+ | Cytidine 5'-diphosphocholine (CDP-choline) | 0.809543269 | 0.270535 |
| 208 | M302T288 | M+ | Arg-Gln | 0.800074723 | 0.110673 |
| 209 | M303T425 | (M+CH3CN+H)+ | Arg-Ser | 0.794540111 | 0.562424 |
| 210 | M173T436_1 | (M-H)- | cis-Aconitate | 0.791416828 | 0.261815 |
| 211 | M88T340_2 | (M-H)- | L-Alanine | 0.772417666 | 0.071754 |
| 212 | M262T450 | (M+H)+ | Lys-Asp | 0.766759792 | 0.274542 |
| 213 | M300T35_2 | (M+H)+ | Palmitoyl ethanolamide | 0.752301013 | 0.284207 |
| 214 | M116T269 | (M-H)- | Acetylglycine | 0.749677481 | 0.282709 |
| 215 | M390T170 | (M+CH3COO+2H)+ | Adenosine 3',5'-cyclic phosphate (cAMP) | 0.74868949 | 0.207182 |
| 216 | M71T400 | (M-H2O-H)- | Dihydroxyacetone | 0.748381183 | 0.038578 |
| 217 | M246T236 | M+ | 2-Methylbutyroylcarnitine | 0.732501894 | 0.150731 |
| 218 | M140T465 | (M-H)- | O-Phosphoethanolamine | 0.727244032 | 0.109809 |
| 219 | M229T373 | (M-H2O-H)- | Pyridoxamine 5'-phosphate | 0.724223173 | 0.541178 |
| 220 | M133T400_2 | (M-H)- | L-Malic acid | 0.713112464 | 0.026855 |
| 221 | M146T372_3 | M+ | (3-Carboxypropyl)trimethylammonium cation | 0.701059885 | 0.005987 |
| 222 | M204T301 | (M+H)+ | Acetylcarnitine | 0.700592895 | 0.423014 |
| 223 | M293T424 | (M+H)+ | EDTA | 0.691865807 | 0.095289 |
| 224 | M284T450 | (M+H)+ | His-Lys | 0.685990806 | 0.116163 |
| 225 | M247T34 | (M+H-2H2O)+ | Oleic acid | 0.674306145 | 0.087024 |
| 226 | M119T34 | (M+H-H2O)+ | Phenylacetic acid | 0.666032471 | 0.003111 |
| 227 | M312T169 | (M+CH3COO+2H)+ | Acetyl Tyrosine Ethyl Ester | 0.660852551 | 0.469769 |
| 228 | M104T370 | (M-H)- | DL-Serine | 0.659536237 | 0.007637 |
| 229 | M364T459 | (M+H)+ | Guanosine 5'-monophosphate (GMP) | 0.656366392 | 0.017106 |
| 230 | M305T35_2 | (M+H)+ | Arachidonic Acid (peroxide free) | 0.649076198 | 0.052275 |
| 231 | M259T347 | (M+H-H2O)+ | 6-Phospho-D-gluconate | 0.644987039 | 0.130621 |
| 232 | M130T270 | (M-H)- | L-Norleucine | 0.635599863 | 0.007446 |
| 233 | M276T399 | (M+H)+ | Arg-Thr | 0.624538322 | 0.178443 |
| 234 | M389T464 | (M+CH3CN+H)+ | 2'-Deoxyguanosine 5'-monophosphate (dGMP) | 0.622115522 | 0.089351 |
| 235 | M662T429 | (M-H)- | Nicotinamide adenine dinucleotide (NAD) | 0.620955762 | 0.074275 |
| 236 | M276T451 | (M+H)+ | .gamma.-L-Glu-.epsilon.-L-Lys | 0.615616583 | 0.109559 |
| 237 | M106T370 | (M+H)+ | L-Serine | 0.603965907 | 0.01342 |
| 238 | M277T54 | (M-H)- | Pantetheine | 0.590005774 | 0.225774 |
| 239 | M179T386_2 | (M-H)- | myo-Inositol | 0.578571427 | 0.260587 |
| 240 | M133T61 | (M+H)+ | Ethyl 3-hydroxybutyrate | 0.572538863 | 0.09312 |
| 241 | M261T356 | (M+H)+ | D-Mannose-6-phosphate | 0.564214504 | 0.099601 |
| 242 | M275T450 | (M-H)- | gamma-L-Glutamyl-L-glutamic acid | 0.560693773 | 0.024156 |
| 243 | M104T148 | M+ | Choline | 0.558721655 | 0.081834 |
| 244 | M338T33_4 | (M+H)+ | Erucamide | 0.557494079 | 0.042253 |
| 245 | M114T167_2 | (M+H)+ | Creatinine | 0.557157756 | 0.492706 |
| 246 | M452T424 | (M-2H+3K)+ | Tyr-Arg | 0.552986137 | 0.002958 |
| 247 | M345T488 | (M+H)+ | Thiamine monophosphate | 0.547757524 | 0.079773 |
| 248 | M116T295_2 | (M-H)- | 5-Aminopentanoic acid | 0.541117068 | 0.008834 |
| 249 | M302T445 | (M+H)+ | N-Acetyl-D-Glucosamine 6-Phosphate | 0.540862358 | 0.039695 |
| 250 | M144T344 | (M+H-H2O)+ | Indole-2-carboxylic acid | 0.516574077 | 0.046831 |
| 251 | M116T483 | (M+H)+ | L-Proline | 0.513423391 | 2.4E-05 |
| 252 | M267T318 | (M+Na-2H)- | Dimethylallyl pyrophosphate | 0.508339333 | 0.049947 |
| 253 | M829T59 | (M+CH3CN+H)+ | N-Docosanoyl-4-sphingenyl-1-O-phosphorylcholine | 0.503435433 | 0.222936 |
| 254 | M244T314_2 | (M+H)+ | Cytidine | 0.493009077 | 0.070142 |
| 255 | M347T443 | (M-H)- | Inosine 5'-monophosphate (IMP) | 0.479934776 | 0.05345 |
| 256 | M787T142 | (M+H)+ | 1,2-dioleoyl-sn-glycero-3-phosphatidylcholine | 0.459363798 | 0.004207 |
| 257 | M147T429 | (M+NH4)+ | L-Pyroglutamic acid | 0.459350009 | 0.000567 |
| 258 | M228T278 | (M+H-2H2O)+ | Gemcitabine | 0.452027259 | 0.007944 |
| 259 | M265T345 | M+ | Thiamine | 0.449150165 | 0.044767 |
| 260 | M115T368 | (M+H-H2O)+ | Ornithine | 0.444392231 | 0.229344 |
| 261 | M284T294 | (M+K)+ | Ile-Asn | 0.443799314 | 0.070779 |
| 262 | M131T414 | (M+H)+ | Agmatine | 0.442615127 | 0.09699 |
| 263 | M204T427 | (M+H-H2O)+ | N-Acetyl-D-glucosamine | 0.441930732 | 3.28E-05 |
| 264 | M116T507 | (M+H)+ | D-Proline | 0.441015795 | 0.059783 |
| 265 | M211T430 | (M-H2O-H)- | D-Ribulose 5-phosphate | 0.414146847 | 0.006985 |
| 266 | M130T519 | (M+H)+ | D-Pipecolinic acid | 0.413656766 | 0.003723 |
| 267 | M147T388_2 | (M-H)- | (S)-2-Hydroxyglutarate | 0.411918981 | 0.012651 |
| 268 | M291T462 | (M+H)+ | Argininosuccinic acid | 0.400480167 | 0.169952 |
| 269 | M187T404 | (M-H)- | N6-Acetyl-L-lysine | 0.386683305 | 0.355777 |
| 270 | M584T439 | (M+NH4)+ | UDP-D-Galactose | 0.386099742 | 9.53E-05 |
| 271 | M219T412 | (M+H)+ | 5-L-Glutamyl-L-alanine | 0.384147043 | 0.007196 |
| 272 | M206T280 | (M+CH3COO+2H)+ | Acetylcholine | 0.375197785 | 0.033455 |
| 273 | M445T441 | (M-H)- | CDP-ethanolamine | 0.366456437 | 0.006235 |
| 274 | M245T409 | (M+H)+ | Pro-Glu | 0.360425775 | 0.001625 |
| 275 | M759T126 | (M+Na)+ | Thioetheramide-PC | 0.354120085 | 0.000407 |
| 276 | M789T142_1 | (M+H)+ | 1-Stearoyl-2-oleoyl-sn-glycerol 3-phosphocholine (SOPC) | 0.349771552 | 0.022931 |
| 277 | M444T449 | (M+H)+ | Guanosine 5'-diphosphate (GDP) | 0.3488213 | 0.004908 |
| 278 | M426T461 | (M-H)- | Adenosine 5'-diphosphate (ADP) | 0.328559662 | 0.00031 |
| 279 | M189T518_3 | (M+H)+ | N6,N6,N6-Trimethyl-L-lysine | 0.323411223 | 0.000207 |
| 280 | M223T454 | (M+H)+ | Allocystathionine | 0.323229412 | 0.010115 |
| 281 | M346T427 | (M-H)- | Adenosine monophosphate (AMP) | 0.318658195 | 0.001136 |
| 282 | M118T346 | (M-H)- | L-Threonine | 0.314666453 | 0.005544 |
| 283 | M90T360 | (M+H)+ | beta-Alanine | 0.310487654 | 0.000485 |
| 284 | M331T34_2 | (M+H)+ | Eicosapentaenoic Acid ethyl ester | 0.30613917 | 0.002531 |
| 285 | M203T485_2 | (M+H)+ | NG,NG-dimethyl-L-arginine(ADMA) | 0.298501815 | 9.68E-05 |
| 286 | M102T365_2 | (M+H)+ | 1-Aminocyclopropanecarboxylic acid | 0.298320673 | 0.00096 |
| 287 | M344T427 | (M+Na)+ | Arg-Phe | 0.292227712 | 0.042 |
| 288 | M795T65 | (M+CH3CN+Na)+ | Sphingomyelin (d18:1/18:0) | 0.285949823 | 0.003337 |
| 289 | M426T424 | (M-H)- | Adenosine 5'-phosphosulfate (APS) | 0.279510219 | 0.007495 |
| 290 | M289T459 | (M+CH3COO)- | D-Ribose 5-phosphate | 0.271260025 | 0.037324 |
| 291 | M606T428 | (M-H)- | UDP-N-acetylglucosamine | 0.269656573 | 1.04E-05 |
| 292 | M148T369 | (M+H)+ | L-Glutamate | 0.263464767 | 0.007318 |
| 293 | M129T388_2 | (M-H)- | Citraconic acid | 0.258926378 | 0.0122 |
| 294 | M565T440 | (M-H)- | Uridine diphosphate glucose(UDP-D-Glucose) | 0.236309608 | 0.000861 |
| 295 | M191T471 | (M-H)- | Citrate | 0.234200667 | 0.014664 |
| 296 | M1005T486 | (M-H)- | Oxytocin | 0.227110351 | 0.009643 |
| 297 | M193T382 | (M-H)- | D-galacturonic acid | 0.223509123 | 0.00596 |
| 298 | M148T278 | (M-H)- | L-Methionine | 0.200275886 | 0.002611 |
| 299 | M400T171_2 | (M+H)+ | L-Palmitoylcarnitine | 0.189843736 | 0.095565 |
| 300 | M156T370 | (M+H)+ | L-Histidine | 0.177536131 | 0.00283 |
| 301 | M74T343 | (M-H)- | Glycine | 0.1752011 | 0.010044 |
| 302 | M428T215 | (2M+NH4)+ | Xanthurenic acid | 0.167325306 | 0.180095 |
| 303 | M464T489 | (M+H)+ | Adenylsuccinic acid | 0.166885587 | 0.001352 |
| 304 | M170T369_2 | (M+H)+ | 3-Methylhistidine | 0.164320912 | 3.69E-05 |
| 305 | M199T445 | (M-H)- | D-Erythrose 4-phosphate | 0.158991184 | 0.002146 |
| 306 | M343T405 | (M+CH3CN+H)+ | Pro-Trp | 0.156024881 | 0.01452 |
| 307 | M611T489 | (M-H)- | Glutathione disulfide | 0.152753417 | 0.015246 |
| 308 | M229T440 | (M+CH3COO)- | Dihydroxyacetone phosphate | 0.148789217 | 0.001778 |
| 309 | M227T410 | (M+CH3COO)- | Phosphoenolpyruvate | 0.145134004 | 0.002022 |
| 310 | M403T427 | (M-H)- | Uridine 5'-diphosphate (UDP) | 0.139036626 | 0.000126 |
| 311 | M145T365_2 | (M-H)- | alpha-ketoglutarate | 0.138975208 | 5.65E-09 |
| 312 | M395T158 | (M-2H+3Na)+ | Tyr-Phe | 0.136300105 | 6.94E-05 |
| 313 | M97T92 | (M+H-H2O)+ | .beta.-Cyano-L-alanine | 0.131559031 | 0.005711 |
| 314 | M76T326 | (M+H)+ | Trimethylamine N-oxide | 0.128671753 | 0.163095 |
| 315 | M261T469 | (M+H)+ | D-Glucose 6-phosphate | 0.122422475 | 0.006746 |
| 316 | M162T348 | (M+H)+ | L-Carnitine | 0.120587404 | 0.013345 |
| 317 | M132T499 | (M+H)+ | Hydroxyproline | 0.119190403 | 0.011523 |
| 318 | M337T461 | (M-H)- | 5'-Phosphoribosyl-5-amino-4-imidazolecarboxamide (AICAR) | 0.098954295 | 0.052556 |
| 319 | M243T468 | (M+H-H2O)+ | alpha-D-Glucose 1-phosphate | 0.096296081 | 0.015867 |
| 320 | M143T339 | (M+CH3CN+H)+ | Betaine aldehyde | 0.071754283 | 0.026594 |
| 321 | M210T162 | (M+CH3COO+2H)+ | Triethanolamine | 0.056413286 | 0.033057 |
| 322 | M259T470 | (M-H)- | D-Mannose 1-phosphate | 0.04892789 | 0.003685 |
| 323 | M124T338 | (M-H)- | Taurine | 0.0390952 | 0.000478 |
| 324 | M110T338_2 | (M+H)+ | Hypotaurine | 0.036079118 | 0.00106 |
| 325 | M230T297_1 | (M+H)+ | Ergothioneine | 0.035366772 | 0.000464 |
| 326 | M259T451 | (M-H)- | alpha-D-Galactose 1-phosphate | 0.035038913 | 0.001011 |
| 327 | M134T143 | (M-H)- | Adenine | 0.012164975 | 0.000231 |
| 328 | M316T189 | M+ | Decanoyl-L-carnitine | 0.000939439 | 0.140775 |
