## Supplementary material for "Integrated multi-approaches reveal unique metabolic mechanisms of Vestimentifera to adapt to deep-sea hydrothermal vents": Table S3

**Table S3. The sequencing data used for the assembly of the *Arcovestia ivanovi* genome.** The coverage was calculated using the estimated genome size with the Kmer-based method. -, not available.

| Library type | Insert size (bp) | Raw data (Gb) | Clean data (Gb) | Read length (bp) | Sequence coverage (X) |
| --- | --- | --- | --- | --- | --- |
| Illumina reads | 350 | 97.8 | 97.5 | 150 | 114.6 |
| Pacbio reads | 8,480 | 94.0 | 94.0 | 7,813 | 110.1 |
| RNA reads | 300 | 9.9 | 9.4 | 150 | - |
