## Supplementary material for "Integrated multi-approaches reveal unique metabolic mechanisms of Vestimentifera to adapt to deep-sea hydrothermal vents": Table S4

**Table S4. The statistics of the** **assembled *Arcovestia ivanovi* genome.**

| Assembly Feature | Statistics |
| --- | --- |
| Number of contigs | 9,469 |
| Total assembly size (bp) | 792,673,544 |
| Longest contig (nt) | 3,857,212 |
| N50 contig length (nt) | 571,012 |
| N90 contig length (nt) | 26,909 |
| BUSCO assessment result | C:97.6%[S:94.9%,D:2.7%], F:2.4%, M:0.0% |
