## Supplementary material for "Integrated multi-approaches reveal unique metabolic mechanisms of Vestimentifera to adapt to deep-sea hydrothermal vents": Table S5

**Table S5. The number of genes in the *Arcovestia ivanovi* genome annotated with different databases.** -, not available.

|  | Number | Percent (%) |
| --- | --- | --- |
| Total | 17,125 | - |
| Swissprot | 13,317 | 77.80 |
| Nr | 15,291 | 89.30 |
| KEGG | 12,706 | 74.20 |
| InterPro | 13,420 | 78.40 |
| GO | 9,770 | 57.10 |
| Pfam | 12,251 | 71.50 |
| Annotated | 15,454 | 90.20 |
| Unannotated | 1,671 | 9.80 |
