## Supplementary material for "Integrated multi-approaches reveal unique metabolic mechanisms of Vestimentifera to adapt to deep-sea hydrothermal vents": Table S6

**Table S6. The categories of the repetitive elements in tubeworms.**

| **Repetitive element category** | ***Lamellibrachia luymesi*** | ***Paraescarpia echinospica*** | ***Arcovestia ivanovi*** | ***Hydroides elegans*** | ***Owenia fusiformis*** |
| --- | --- | --- | --- | --- | --- |
| DNA | 252849 | 532512 | 194359 | 103732 | 1319730 |
| DNA/Academ-1 | 3344864 | 15410928 | 6264981 | 168759 | 1789824 |
| DNA/Academ-2 | 272941 |  |  |  | 8879 |
| DNA/CMC-Chapaev | 51345 |  |  |  | 69869 |
| DNA/CMC-Chapaev-3 |  |  | 138786 |  |  |
| DNA/CMC-EnSpm | 294147 | 1096683 | 210182 | 707410 | 59364 |
| DNA/Crypton |  |  |  | 427249 | 44828 |
| DNA/Crypton-A |  | 186783 |  | 308150 | 83839 |
| DNA/Crypton-V | 109289 | 332499 | 174297 | 76372 |  |
| DNA/Dada |  | 258068 |  |  | 105236 |
| DNA/Ginger-1 | 19740 | 138937 | 236790 |  | 19330 |
| DNA/Ginger-2 | 88893 |  |  | 138782 |  |
| DNA/hAT | 158534 | 468830 | 168462 | 53467 |  |
| DNA/hAT-Ac | 918668 | 2107587 | 1521591 | 100948 | 65415 |
| DNA/hAT-Blackjack | 441869 | 974414 | 340811 |  |  |
| DNA/hAT-Charlie | 1274097 | 3265709 | 1690639 |  | 111054 |
| DNA/hAT-hAT5 | 236422 | 482663 | 247597 | 406613 | 22121 |
| DNA/hAT-hAT6 | 11643 | 132094 | 14544 |  | 219201 |
| DNA/hAT-hATm | 274863 | 389362 | 27047 |  | 76957 |
| DNA/hAT-hATw | 40565 | 335606 |  |  |  |
| DNA/hAT-hobo |  |  |  |  | 54157 |
| DNA/hAT-Tag1 | 791209 | 276304 | 304620 |  |  |
| DNA/hAT-Tip100 | 2850676 | 4647710 | 1651348 | 159562 | 2110325 |
| DNA/IS3EU | 82263 | 898061 | 96102 |  | 77668 |
| DNA/Kolobok-Hydra | 497651 | 822135 | 736704 |  |  |
| DNA/Kolobok-T2 | 404135 | 2935208 | 1205479 | 45190 | 97435 |
| DNA/Maverick | 2699253 | 34188628 | 6787715 | 3382426 | 686554 |
| DNA/Merlin | 132712 | 120519 | 83324 |  |  |
| DNA/MULE | 27552 |  |  |  |  |
| DNA/MULE-MuDR | 570189 | 1173189 | 226258 | 389088 | 730856 |
| DNA/MULE-NOF | 43043 | 93545 |  |  |  |
| DNA/P | 838947 | 1468302 | 289744 | 195165 | 33649 |
| DNA/PIF-Harbinger | 253409 | 811922 | 74258 | 272469 | 238555 |
| DNA/PIF-ISL2EU | 258371 | 1203903 | 240095 | 199562 |  |
| DNA/PIF-Spy |  |  |  |  | 132845 |
| DNA/PiggyBac | 281052 | 1560041 | 470469 |  | 22782 |
| DNA/Sola-1 | 329701 | 313036 | 352923 | 48177 |  |
| DNA/Sola-2 | 976785 | 2200642 | 1406761 |  | 82240 |
| DNA/TcMar-Fot1 | 1304233 | 2927268 | 643302 | 300005 |  |
| DNA/TcMar-m44 |  |  |  | 931657 |  |
| DNA/TcMar-Mariner |  | 173879 | 30585 | 110578 | 6009516 |
| DNA/TcMar-Pogo | 304173 | 272959 | 104536 | 262519 |  |
| DNA/TcMar-Tc1 | 621599 | 882686 | 409597 | 4698400 | 3966508 |
| DNA/TcMar-Tc2 | 139651 |  |  |  |  |
| DNA/TcMar-Tigger | 1329126 | 945988 | 83147 | 948274 |  |
| DNA/Zator | 42230 |  |  |  |  |
| LINE | 442248 | 3062825 | 640671 | 1771628 | 992138 |
| LINE/CR1 | 36194580 | 73464829 | 62675865 | 7050715 | 677463 |
| LINE/CR1-Zenon | 406102 | 245207 | 158524 |  | 2232709 |
| LINE/I |  | 103176 |  | 162374 | 102946 |
| LINE/I-Jockey | 897298 | 910950 | 678693 | 10716 | 266744 |
| LINE/L1 |  | 573587 | 46391 | 116842 | 46594 |
| LINE/L1-Tx1 | 453409 | 1218737 | 355955 | 5999603 | 2401949 |
| LINE/L2 | 20685310 | 61545588 | 29773092 | 11829635 | 3002585 |
| LINE/Penelope | 1974097 | 4065250 | 1685971 | 408031 | 990703 |
| LINE/Proto2 | 3407750 | 5982474 | 1953265 | 484782 | 638061 |
| LINE/R1 |  |  | 141343 |  | 1098 |
| LINE/R1-LOA |  |  | 40019 |  |  |
| LINE/R2 | 157544 | 522834 | 338669 |  | 19389 |
| LINE/R2-Hero |  | 554174 | 156563 | 5876817 | 533442 |
| LINE/R2-NeSL |  | 33026 |  |  |  |
| LINE/Rex-Babar | 2095489 | 5493590 | 2915512 | 1056380 |  |
| LINE/RTE | 944378 | 1310065 | 2052843 | 1472515 | 258683 |
| LINE/RTE-BovB | 17742421 | 31035538 | 15083189 | 6818729 |  |
| LINE/RTE-RTE | 748489 | 4581138 | 4832514 |  |  |
| LINE/RTE-X | 800648 | 10845178 | 518826 | 972987 | 2768772 |
| LINE/Tad1 |  |  |  |  | 36724 |
| Low_complexity | 614812 | 877764 | 1044036 | 1487704 | 380986 |
| LTR |  |  | 15983 | 110358 |  |
| LTR/Copia | 170094 | 17747 |  |  |  |
| LTR/DIRS | 60691 | 410024 | 176281 | 3971397 | 803153 |
| LTR/ERV-Foamy |  | 95735 |  |  |  |
| LTR/ERV1 |  | 27463 | 49413 |  |  |
| LTR/ERVL | 65763 | 14376 |  |  |  |
| LTR/Gypsy | 5507643 | 28750587 | 3715011 | 5735829 | 4221566 |
| LTR/Ngaro | 1442419 | 3784290 | 2407628 | 444041 | 994801 |
| LTR/Pao | 306071 | 687100 | 173711 | 2191164 |  |
| RC/Helitron | 101726 | 506114 | 150592 | 2434959 | 1580725 |
| rRNA |  | 669818 | 1360877 | 287657 | 55285 |
| Satellite | 663426 | 825473 | 35422 | 363019 | 102212 |
| Simple_repeat | 10888615 | 28292981 | 36505670 | 9236203 | 3023268 |
| SINE |  |  |  |  | 103282 |
| SINE? | 3235 |  |  |  | 4777 |
| SINE/B4 |  |  |  | 255458 |  |
| SINE/MIR | 4706250 | 9250668 | 6253147 |  |  |
| SINE/tRNA |  |  |  |  | 420390 |
| SINE/tRNA-V-RTE |  |  |  | 379924 |  |
| snRNA |  | 20364 |  | 33687 |  |
| tRNA | 398562 | 1986438 | 24732 | 4994449 | 607850 |
| Unknown | 168168880 | 338969019 | 275052197 | 308461597 | 215242413 |
