## Supplementary material for "Integrated multi-approaches reveal unique metabolic mechanisms of Vestimentifera to adapt to deep-sea hydrothermal vents": Table S7

**Table S7. The central carbon metabolisms in Polychaeta.**

|  | Glycolysis | Tricarboxylic acid cycle | Pentose phosphate pathway | Glyconeogenesis | Trehaloneogenesis | Glycogenesis |
| --- | --- | --- | --- | --- | --- | --- |
| **Symbiotic Polychaeta** |  |  |  |  |  |  |
| **Vestimentifera** |  |  |  |  |  |  |
| *Arcovestia ivanovi* | + | + | + | - | + | + |
| *Oasisia alvinae* | + | + | + | - | + | + |
| *Riftia pachyptila* | + | + | + | - | + | + |
| *Paraescarpia echinospica* | + | + | + | - | + | + |
| *Lamellibrachia luymesi* | + | + | + | - | + | + |
| **Osedax** |  |  |  |  |  |  |
| *Osedax frankpressi* | + | + | + | - | - | + |
| **Non symbiotic Polychaeta** |  |  |  |  |  |  |
| *Capitella teleta* | + | + | + | + | - | + |
| *Dimorphilus gyrociliatus* | + | + | + | + | - | + |
| *Owenia fusiformis* | + | + | + | + | - | + |
| *Hydroides elegans* | + | + | + | + | - | + |

“+” and “–” indicate presence and absence, respectively, of the specified pathway.
